# Site-specific Covalent Labeling of DNA Substrates by an RNA Transglycosylase

**DOI:** 10.1101/2023.01.23.525207

**Authors:** Ember M. Tota, Neal K. Devaraj

**Affiliations:** Department of Chemistry and Biochemistry, University of California, San Diego, 9500 Gilman Drive, La Jolla, California 92093, United States

**Author notes:** Corresponding Author Neal K. Devaraj − Department of Chemistry and Biochemistry, University of California, San Diego, La Jolla, California 92093, United States.

## Abstract

Bacterial tRNA guanine transglycosylases (TGTs) catalyze the exchange of guanine for the 7-deazaguanine queuine precursor, prequeuosine1 (preQ1). While the native nucleic acid substrate for bacterial TGTs is the anticodon loop of queuine-cognate tRNAs, the minimum recognition sequence for the enzyme is a structured hairpin containing the target G nucleobase in a “UGU” loop motif. Previous work has established an RNA modification system, RNA-TAG, in which E. coli TGT exchanges the target G on an RNA of interest for chemically modified preQ1 substrates linked to a small molecule reporter such as biotin or a fluorophore. While extending the substrate scope of RNA transglycosylases to include DNA would enable numerous applications, it has been previously reported that TGT is incapable of modifying native DNA. Here we demonstrate that TGT can in fact recognize and label specific DNA substrates. Through iterative testing of rationally mutated DNA hairpin sequences, we determined the minimal sequence requirements for transglycosylation of unmodified DNA by E. coli TGT. Controlling steric constraint in the DNA hairpin dramatically affects labeling efficiency, and, when optimized, can lead to near quantitative site-specific modification. We demonstrate the utility of our newly developed DNA-TAG system by rapidly synthesizing probes for fluorescent Northern blotting of spliceosomal U6 RNA and RNA FISH visualization of the long noncoding RNA, MALAT1. The ease and convenience of the DNA-TAG system will provide researchers with a tool for accessing a wide variety of affordable modified DNA substrates.

## Introduction

Probe modified oligonucleotides have widespread applications in imaging, diagnostics, nanotechnology, and medicine.^1–3^ Modified nucleic acids are commonly produced by incorporating functionalized nucleotides during solid-phase synthesis using phosphoramidite chemistry. In addition, several alternative techniques exist that rely on chemical or enzymatic modification of nucleobases or the sugar backbone.^4–11^ Techniques that employ enzymes offer several advantages over those relying on chemical modification as they occur in mild and aqueous conditions, are precisely targeted, and are typically very efficient. The deoxyribose backbone of DNA affords aqueous stability that is lacking in its RNA counterpart, and as such, chemically functionalized single-stranded DNAs (ssDNA) are widely used in biotechnology. Current methods for enzymatic modification of DNA include the 3′ insertion of modified nucleobases using terminal deoxynucleotidyltransferase (TdT) and the 5′ insertion of modified phosphate groups.^12,13^ Unfortunately, the small molecule substrate tolerance and precision of insertion for enzyme-mediated ssDNA modification strategies has been limited. The improved ability to site-specifically label ssDNA with enzymes in mild aqueous conditions could have significant applications given the broad use of ssDNA in diagnostics, therapeutics, materials research, imaging, detection, and chemical barcoding.^11,14–16^

Previously, our group developed RNA-TAG (RNA **t**ransglycosylation **a**t **g**uanosine) to site-specifically label RNA using bacterial tRNA guanine transglycosylase (TGT).^17,18^ TGTs catalyze the exchange of guanine for 7-deazaguanine derivatives on the anticodon loop of queuine-cognate tRNAs.^19^ These enzymatic modifications are present across all three kingdoms of life, differing slightly in their small molecule and tRNA substrate counterparts. It had previously been shown that the entire tRNA structure is not necessary for enzymatic activity of *E. coli* TGT toward RNA substrates, but rather, the minimum recognition sequence is a short 17-nucleotide hairpin (ECY-A1; the anticodon loop of tRNA^Tyr^) which contains a uridine-flanked target guanine in its loop.^20^ In RNA-TAG, *E. coli* TGT recognizes this minimal hairpin incorporated into an RNA of interest and catalyzes the exchange of the target guanine for synthetic derivatives of the natural preQ1 substrate that are conjugated to functional groups such as fluorophores or affinity ligands.^17^ We have demonstrated the utility of RNA-TAG in a range of applications including RNA biotinylation for proteomics studies, controlling mRNA translation, and light activated CRISPR gene editing in cells.^17,18,21–26^

Extending the RNA-TAG system to work on DNA would enable one-step, site-specific labeling of ssDNA substrates, increasing the accessibility of small molecule modified DNA oligonucleotides and further enabling applications of ssDNA in nanotechnology, therapeutics, and basic research. The exchange of guanine for preQ1 by TGT is electronically dependent on the guanine nucleobase and the endocyclic oxygen of the ribose backbone.^19^ This suggests that the lack of the exocyclic 2’ hydroxyl groups on the backbone of DNA substrates would not impair *E. coli* TGT recognition or activity. However, previous work had deemed *E. coli* TGT incapable of acting on unmodified DNA substrates and showed that unnatural nucleobase substitutions (T→dU) of the thermally stabilized 4 base-pair extended-stem mini helix (dECYMH) were required for efficient recognition.^19,27^

We speculated that optimizing the hairpin structure could enable recognition of native DNA by *E. coli* TGT. Indeed, through iterative testing of rationally designed DNA hairpin sequences, we were able to reveal the minimal sequence requirements for DNA substrate recognition by the enzyme (Figures 1A and S3), enabling development of a technique we term DNA **t**ransglycosylation **a**t deoxy**g**uanosine, or DNA-TAG. DNA-TAG enables the one-step, site-specific labeling of ssDNA substrates bearing a minimal 17-nucleotide hairpin motif with synthetic preQ1 substrates conjugated to functional groups such as biotin and fluorophores (Figure 1B). Next, we characterize the ability of DNA-TAG to label DNAs that include one or more recognition sequences and explore the general substrate scope for DNA-TAG. Finally, we show how DNA-TAG can be used to generate probes quickly and inexpensively for fluorescent Northern blotting and RNA FISH.

**Figure 1.**
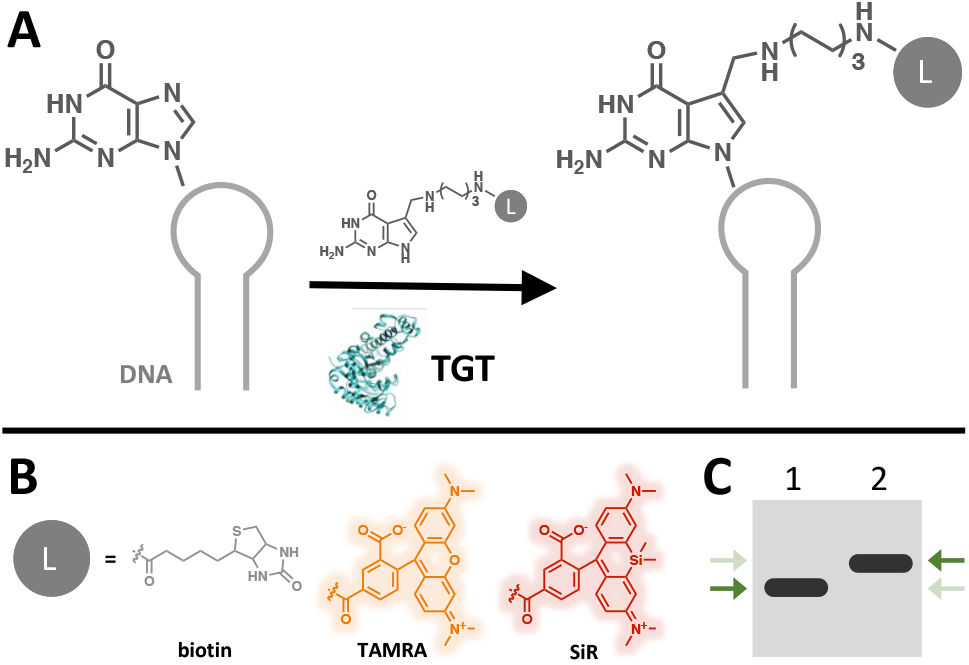
Proposed scheme of *E. coli* TGT mediated modification of DNA with functional small molecules. (A) General scheme of DNA-TAG. *E. coli* TGT irreversibly exchanges a specific guanine with a preQ1-ligand (L) on a DNA hairpin. (B) PreQ1-ligands used in this work include preQ1-biotin, preQ1-tetramethyl-rhodamine (TAMRA), and preQ1-silicon rhodamine (SiR). (C) Cartoon depiction of expected upward gel shift due to increased mass from the base exchange reaction as indicated by the green arrows. Lane 1: DNA; Lane 2: preQ1-L modified DNA.

## Results and Discussion

Previous studies reported that TGT does not modify the DNA equivalent substrate (dECYMH) of the extended stem tRNA^tyr^-based RNA mini helix (ECYMH). However, it was also reported that when all “T” nucleobases in the hairpin were replaced with “U”, the enzyme was able to recognize and form the covalent intermediate with a deoxyribose-based substrate (dUdECYMH).^27^ While we confirmed only trace modification (less than 5% by densitometry) of dECYMH with our preQ1-biotin probe (Figure 2A-B), the published mechanism of TGT catalysis alludes to the possibility that the enzyme could recognize unmodified DNA substrates.^19^ With this information, we designed a series of experiments to assess the limitations of TGT toward accepting DNA, restricting the number of T→dU mutations to only those that are part of the minimal “UGU” recognition element in the extended stem DNA analog dECYMH: <u>GGGA</u>*GCAGA<u>C**XGX**AAA</u>TCTGC*<u>TCCC</u> (dECY-A1 is italicized, the 4 base-pair extension and loop sequences are underlined, and the minimum recognition element is bolded). For all DNA substrate modification experiments, we relied on an established oligonucleotide gel shift assay.^18,23,26^ Observation of an upward gel shift of the product in relation to the starting material reports an increase in mass from the enzymatic exchange of guanine for a preQ1-biotin small molecule reporter conjugate (Figures 1B-C and 2A). We surveyed transglycosylase activity toward dECYMH hairpin variants containing the following minimal recognition element identities: TGT, dUGdU, dUGT, and TGdU. While the TGT and dUGT containing hairpins showed no appreciable modification by the enzyme, those containing dUGdU and TGdU were modified to a similar extent as the wildtype RNA substrate, as evidenced by the upward gel shift in the product lane (Figure 2B). It appears that a single nucleobase mutation of the minimal recognition element from TGT to TGdU is sufficient for TGT to recognize and modify the dECYMH substrate. This finding expands on the previous observation that turnover of a DNA substrate only occurred when all T nucleotides in the hairpin were replaced with dU.^27^ Encouraged by this result, we next set out to ask which aspects of dECYMH were limiting TGT enzyme activity. We synthesized an RNA hairpin (rECYMH-rTGrT) in which all U residues were replaced with 5-meU (rT). Surprisingly, we found that this substrate had a nearly quantitative turnover, similar to the wildtype RNA hairpin (Figure 2C), indicating that TGT can efficiently modify nucleic acid substrates containing thymine bases. Combined with the fact that a minimally modified DNA sequence with a single T→dU mutation is also recognized, our initial findings suggested that the inability of *E. coli* TGT to modify dECYMH likely arises from the combined steric constraints of the deoxyribose backbone and the methyl groups on the T bases of the putative dTdGdT recognition element.

**Figure 2.**
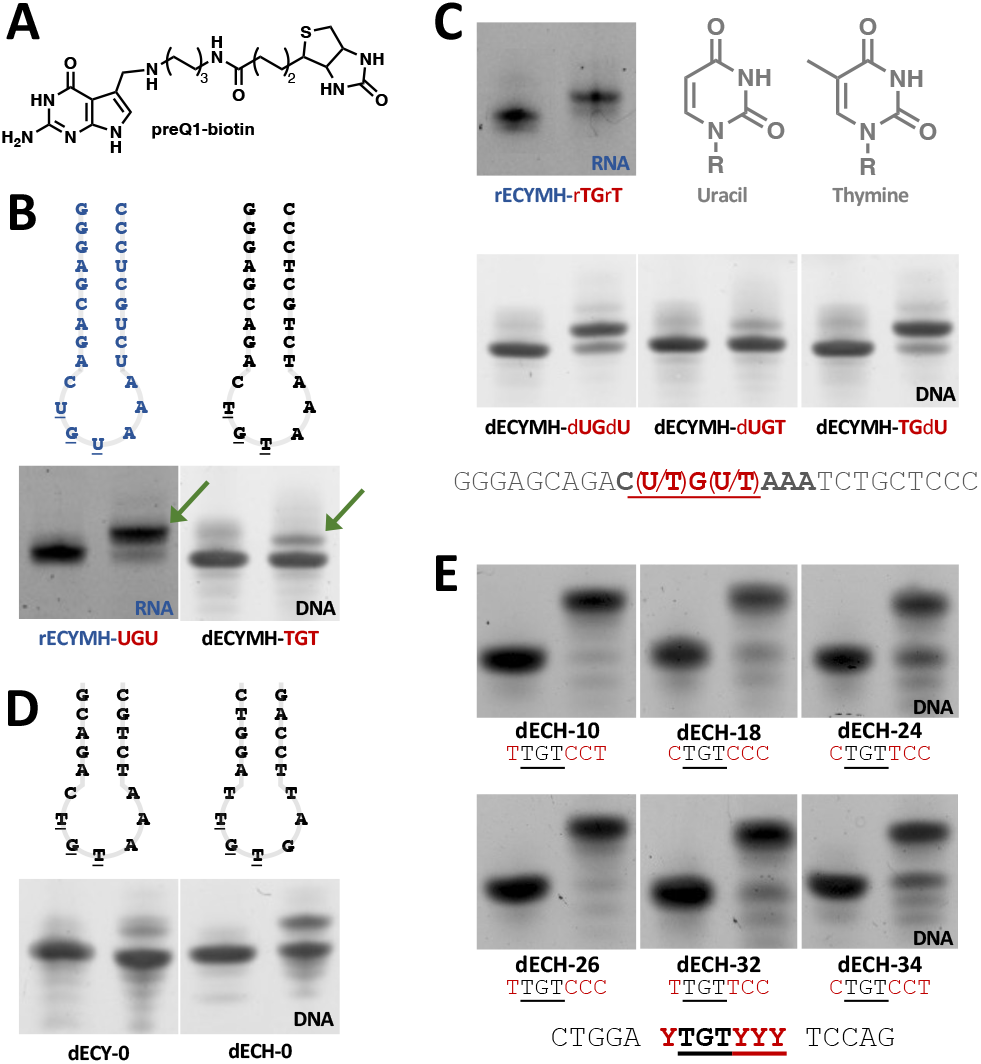
Determining the minimal sequence requirements for DNA modification by *E. coli* TGT. For gel data, unmodified oligos are in the left lane and reaction products are in the right lane. Modification is determined by an upward gel shift of the oligo after insertion of preQ1-biotin. (A) PreQ1-biotin probe. (B) TGT modification of extended stem RNA and DNA hairpins derived from the anticodon loop of tRNA^tyr^; arrows indicate modified product as is evidenced by the upward gel shift. (C) TGT labeling of T→dU mutants. A single mutation is sufficient for TGT modification of dECYMH. (D) TGT modifies dECY-0 and dECH-0 at ~5% and ~40% efficiency, respectively. (E) TGT modifies DNA-based hairpins with >90% efficiency when the loop sequence is Y<u>TGT</u>YCC or Y<u>TGT</u>CCY where Y is C or T, except for C<u>TGT</u>TCC, which is labeled with ~85% efficiency.

Based on these observations, we hypothesized that, rather than a fundamental incompatibility of TGT with DNA oligonucleotides, the seeming inability of TGT to modify DNA was steric and could be overcome by using an alternative DNA substrate. Previous studies have focused on nucleic acid substrates related to the 17-nucleotide ECY-A1 and 25-nucleotide ECYMH hairpins which are derived from the anticodon loop of the tyrosine tRNA containing the GUA anticodon. However, all four tRNAs with GUN anticodons are substrates for TGT, including tRNA^Tyr^, tRNA^His^, tRNA^Asp^, and tRNA^Asn^. While each of these four tRNAs have the same “UGU” minimal recognition element, the rest of the anticodon loop sequence varies. Therefore, we decided that hairpin substrates based on these alternative tRNAs would be a good starting point for engineering a novel DNA-TAG substrate. Adapting the already established nomenclature, and to account for mutations, we refer to the 17-nucleotide wildtype hairpins as dECY-0 (originally dECY-A1), dECH-0, dECD-0, and dECN-0, corresponding to the hairpins derived from tRNA^Tyr^, tRNA^His^, tRNA^Asp^, and tRNA^Asn^, respectively. Mutant hairpins are referred to using a sequential numbering scheme based on their parent hairpin (e.g., dECY-1, dECY-2, etc.). All hairpins were treated with *E. coli* TGT and preQ1-biotin and the resulting products analyzed via urea PAGE gel shift. Interestingly, both the dECH-0 and dECD-0, hairpins showed an appreciable increase in TGT labeling activity compared to their dECY-0 counterpart, with labeled product increasing from ~5% to ~40% by densitometry (Figures 2D and S1). We note that dECN-0 shows a shift indicative of degradation rather than modification, a phenomenon that is consistent for most oligos bearing a 3′ T (Figures S1, S2, and S4). While the data was included here for completeness, it is outside the scope of this work and will not be discussed further. The approximate increase in labeling of the dECH-0 and dECD-0 substrates to 40% was promising, but not comparable to the near quantitative labeling observed with RNA-TAG.^17^ To further improve the DNA-TAG substrate, we next sought to determine the importance of the stems and/or loops for each of the tRNA derived hairpins. We designed composite substrates, swapping the stems and loops of all four cognate DNA analogs. All hairpins with the aspartate-based loop (dECD) demonstrated labeling of over 40%, with the histidine stem-aspartate loop substrate reaching over 50% by densitometry (dECH-dECD; Figure S1). The results of this experiment suggested that while both the stem and the loop sequences contribute to the ability of TGT to turnover a given substrate, the loop sequence plays the more critical role in substrate compatibility, in line with previous observations during biochemical and structural studies of the enzyme and tRNA substrate.^20,28^

We decided to move forward with the dECH-0 substrate for further development and designed 9 hairpin mutants (dECH-1-9; Figure S2). These experiments confirmed the importance of the loop and led us to design and test another 31 mutants (dECH-10-40; Figure S3). This round of mutants produced optimal DNA-TAG substrates and revealed the minimal substrate loop requirement for efficient recognition and modification of DNA by TGT to be either YTGTCCY or YTGTYCC, where Y represents smaller pyrimidine bases (C or T), further corroborating our hypothesis that steric effects play a major role in substrate recognition (Figure 2E). We chose one of the hairpins with the highest labeling efficiency, dECH-10, as our model substrate to probe how stem sequence affects DNA modification by TGT. Stems derived from the anticodon arm sequences of all *E. coli* tRNAs were appended with the dECH-10 “TTGTCCT” loop (Figure S4). Nearly all the anticodon-based hairpins were appreciably labeled regardless of the stem sequence, apart from some apparent multi-labeling and degradation products, similar to that seen for dECN-0. Ultimately, our data suggests that, with the right loop, the stem is far less important, as 16 DNA hairpins with different stem sequences are nearly quantitatively labeled by *E. coli* TGT (all greater than 95% by densitometry; Figure S5).

While any of the 16 substrates are valid candidates for use with DNA-TAG, we performed subsequent experiments with dECH-10 (hairpin 176, CTGGA**<u>TTGTCCT</u>**TCCAG). To confirm the result of our gel shift assay, we verified labeling with liquid chromatography-mass spectrometry analysis (LCMS). The LC trace showed a clear product peak with the expected mass, matching the conclusion derived from urea PAGE gel shift analysis (Figure S6). We verified that the base being exchanged by the enzyme was the expected G residue by testing whether dECH-10ΔC (CTGGA<u>TT**C**TCCT</u>TCCAG) could be modified with TGT and found that the mutant hairpin remained unlabeled (Figure S7).

Having optimized DNA hairpin labeling by TGT, we sought to explore the utility of our newly developed DNA-TAG system. A key benefit of RNA-TAG is the wide substrate tolerance of the enzyme for preQ1 derivatives. We compared the substrate tolerance for DNA modification and found that, similar to RNA, TGT can modify the dECH-10 substrate with a variety of functional preQ1 probes in high yield. We used the established gel shift assay to assess TGT’s ability to modify dECH-10 with various probes: preQ1 – biotin, TAMRA, Cy5, Alexa Fluor 647, Alexa Fluor 488, and silicon rhodamine (Figure S8) and confirmed with high resolution mass spectrometry (HRMS; see supporting information).

DNA-TAG modification requires that a TGT recognition harpin be appended to the DNA of interest. We purchased 59-mer DNA constructs containing 5′, 3′, and internal hairpins and tested the extent of their labeling. All three hairpin adducts were efficiently labeled using the preQ1-biotin probe (Figure S9). Additionally, we tested the ability of DNA-TAG to insert multiple labels into a single DNA oligonucleotide. Indeed, TGT modification of transcripts bearing two hairpins either in tandem at the same end (5′, 5′ or 3′, 3′) or separated at opposite ends (5′, 3′) was observed via gel shift and confirmed with LC-ESI-TOF-MS (Figure S10). Our experiments demonstrate that DNA-TAG has the potential to be used in a range of applications as it is compatible with several small molecule probes varying in size and overall charge. Furthermore, DNA-TAG can be used for labeling DNA oligonucleotides at a location of choice, both singly and in tandem.

DNA-TAG could offer a simple, efficient, and cost-effective means for generating fluorescent probes to reliably detect nucleic acid species of interest. We initially explored the suitability of DNA-TAG for generating fluorescent probes for Northern blotting. Northern blotting is a technique used to visualize or quantify an RNA species from a given sample. In short, the sample is separated by gel electrophoresis, transferred to a membrane, and detected via hybridization of reporter labeled antisense oligonucleotide probes. Conventional Northern blotting utilizes radiolabeled probes because they are highly sensitive, and the methods are well established. However, radioactive probes require significant safety measures due to their toxicity and only produce a single output signal, restricting multiplexing capabilities. Recently developed fluorescent blotting strategies, on the other hand, have many advantages over radiolabeling including mitigating safety concerns and allowing for detection of different targets via the use of orthogonal fluorophores.^29^ We designed a single fluorescently labeled probe to detect U6 RNA, a small nuclear RNA which plays a catalytic role in the spliceosome.^30^ A 50-mer DNA oligonucleotide U6 antisense probe was appended with a 5′ DNA-TAG hairpin (Figure 3A) and labeled with preQ1-silicon rhodamine (preQ1-SiR; Figure 1B), a near infrared fluorophore used in cell imaging.^31^ Samples containing *in vitro* transcribed (IVT) U6 RNA or crude total RNA from U2OS cellular extracts were separated via urea PAGE in dilution series. The gels were transferred to a nitrocellulose membrane, incubated with the SiR labeled antisense probe, and the signal detected via fluorescence scanning. The DNA-TAG generated U6 Northern blot probe was successful in detecting the U6 RNA in a dose dependent manner with a reliable R^2^ value when analyzed via linear fit for both IVT and cellular U6 RNA (Figure 3B-C, respectively). The SiR fluorescent Northern blot detection appears to be comparable to previously reported detection limits (Figure 3B).^29,32^ In the future, the sensitivity and specificity of the technique might be improved by using reporter fluorophores with better photophysical properties, including multiple fluorescent tags on a single probe, or generating multiple probes and using a tiled detection approach.

**Figure 3.**
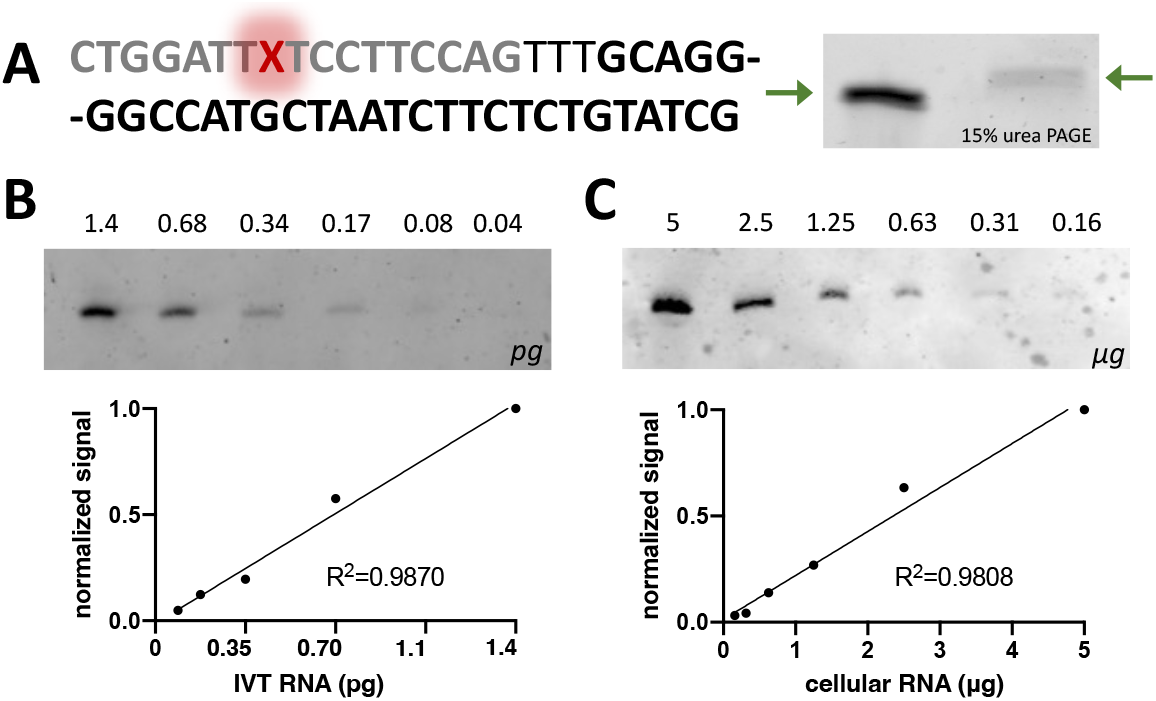
Near-infrared fluorescent Northern blot of U6 RNA using a DNA-TAG generated SiR-antisense probe. (A) SiR-labeled antisense probe. The U6 antisense sequence is black; the DNA-TAG recognition sequence is grey; the inserted preQ1-SiR (X) probe is red. Modification of the anti-sense probe with preQ1-SiR is evidenced by an upward gel shift; unmodified antisense probe is on the left and SiR labeled anti-sense probe is on the right; note that the gel red signal is quenched by the SiR dye. (B) Dose dependent fluorescence detection of picogram (pg) quantities of in vitro transcribed U6 RNA. (C) Dose dependent fluorescence detection of U6 RNA from microgram (μg) quantities of total cellular RNA extract from U2OS cells.

Next, we used DNA-TAG to generate probe sets for single molecule FISH (smFISH), a powerful technique that has been widely used to visualize specific DNA and RNA species in cells using tiled antisense probes labeled with fluorescent reporters.^33–36^ There are a variety of smFISH techniques, all of which rely on the use of fluorescently labeled nucleic acid probes. FISH probes have been generated using naturally occurring nucleic acids such as DNA and RNA, and synthetic peptide nucleic acids (PNA).^37^ Imaging signals can be increased with a variety of strategies, including increasing the number of probes or reporter molecules^38^ and using turn-on intercalating dyes, such as thiazole orange, to minimize background.^16^ A standard smFISH probe set is made up of 30-48 distinct probes which tile the target transcript.^39^ Given the large number of probes required, probe set synthesis can be very costly and needs to be high-yielding. While labeled probe sets can be purchased commercially or generated using TdT along with a limited set of accepted ddUTP-fluorophores, DNA-TAG offers a unique strategy for quickly generating affordable probe sets with a wide small molecule substrate scope.^40^ We tested the ability of DNA-TAG to generate a fluorescent-tiled antisense oligo probe set against MALAT1.

MALAT1 (metastasis-associated lung adenocarcinoma transcript 1) is an abundant long non-coding RNA (lncRNA) found in the nuclei of cells that is implicated in cancers including those affecting the lungs, breasts, prostate, pancreas, glia, and blood.^41^ A set of 48 unique antisense oligonucleotide probe sequences was selected to target MALAT1 and appended with DNA-TAG hairpins. We designed two distinct probe sets containing either one or two TGT modification sites. To maintain consistency between the sets, both constructs had two hairpins, with the single modification set containing one hairpin bearing the “TGT” minimal recognition element and the other containing the inactive “TCT” minimal recognition element (Tables S7 and S8). The probes were individually purchased from a commercial vendor (Integrated DNA Technologies), pooled, and fluorescently labeled using TGT and preQ1-TAMRA. The combined oligonucleotide probe set was then purified by spin filtration. U2OS cells were treated with either a DNA-TAG-labeled TAMRA probe set, an orthogonal Quasar 570-labeled commercial probe set (Stellaris), or a combination of the two. As expected for MALAT1, treatment with all probe sets resulted in punctate stains localized in the nuclei of the cells, with both DNA-TAG labeled probe sets colocalizing with the Stellaris probe set (Figure 4). The signal from the single label probe set was sufficient for reliably detecting MALAT1 and we did not observe significant improvement in signal from doubly labeled probes (Figures S11 and S12). The latter observation may be due to the well-known self-quenching of TAMRA and further optimization of fluorophore spacing may result in improved signals.^42^ A single experimental design was shown here, however, our strategy should be amenable with a variety of amplification strategies, for instance, by optimizing spacing and including additional hairpins on each probe to increase the fluorescence signal.

**Figure 4.**
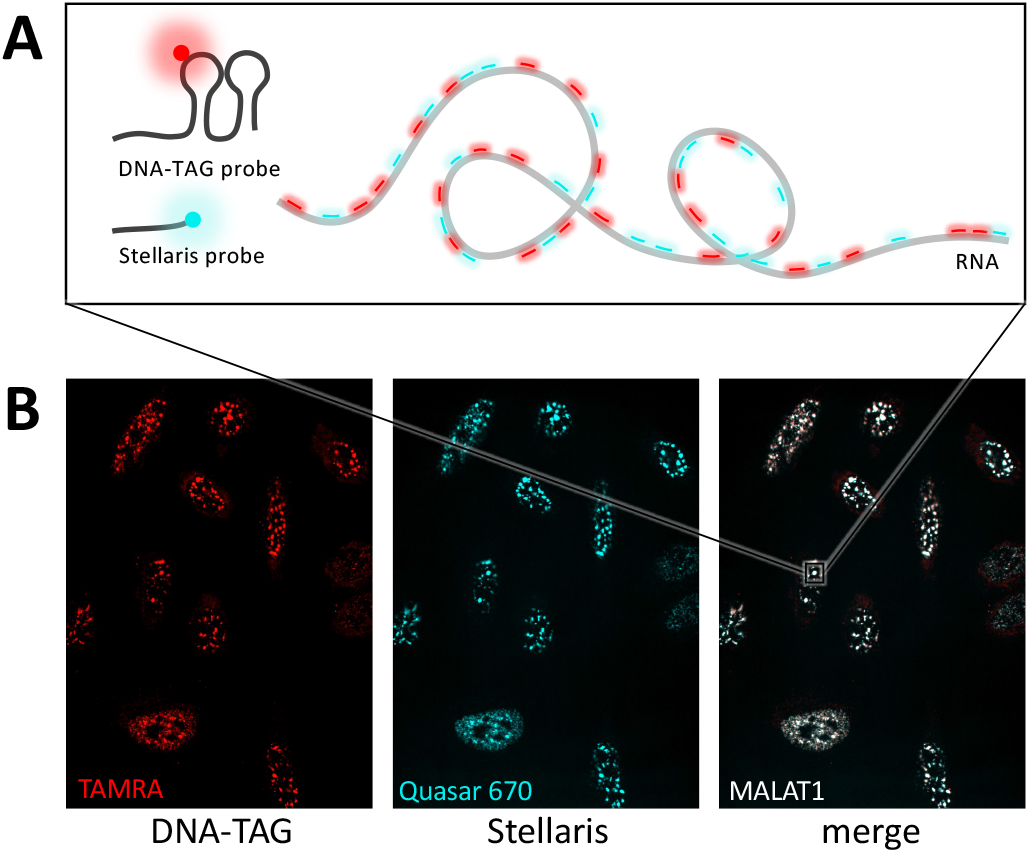
RNA-FISH detection of MALAT1 RNA in U2OS cells using both a DNA-TAG-generated probe set and a commercial probe set from Stellaris. (A) Cartoon depiction of RNA-FISH using the tiled probe sets. The DNA-TAG probe set is modified with preQ1-TAMRA (red) and the Stellaris probe set is labeled with Quasar 670 (teal). (B) Both probe sets hybridize with the MALAT1 puncta in the nuclei of U2OS cells.

## Conclusion

Beyond being central to life, nucleic acids have proven to be invaluable tools in both basic and biomedical research. Harnessing the unparalleled programmability afforded by oligonucleotides requires tools to install functional handles or facilitate detection. Furthermore, increasing the availability, versatility, and diversity of oligonucleotide tools is critical to the continued advancement of their use in multiple fields. We have shown that a bacterial TGT enzyme, which normally modifies tRNA, can be repurposed to site-specifically label DNA. Enzymatic modification using the DNA-TAG methodology facilitates the one-step, site-specific insertion of a variety of functional small molecules into ssDNA substrates of interest for downstream applications. This system is compatible with both internal and terminal insertions of the hairpin sequence into DNAs of interest, is tolerant of tandem modifications, and works with a variety of small molecule substrates. Furthermore, DNA-TAG allows an inexpensive and straightforward method by which researchers can quickly label several DNA oligos in a single step, either simultaneously or in parallel, followed by a short spin column purification. We have demonstrated that, with a one-time upfront cost of materials needed for small molecule probe synthesis and protein expression, DNA-TAG offers researchers a versatile and affordable means to generate functional or fluorescent DNA probes of their choice. While we have demonstrated applications such as fluorescence Northern blotting and smFISH, given the modularity of the approach, numerous techniques and technologies that rely on modified oligonucleotides may benefit from DNA-TAG probe synthesis.

## Supporting information

Tota_Devaraj_Supplemental Information

## Acknowledgements

The authors acknowledge the UCSD Molecular Mass Spectrometry Facility for small molecule high resolution and oligonucleotide mass spectrometry support. This work was supported by the National Institutes of Health [R35 GM141939]. The content of the information does not necessarily reflect the position or the policy of the government, and no official endorsement should be inferred.

