## Supplementary material for "Site-specific Covalent Labeling of DNA Substrates by an RNA Transglycosylase": Tota_Devaraj_Supplemental Information

**Table of Contents**

1. **General Information**
2. **Buffers**
3. **Plasmid Design and Sequences**
4. **Oligonucleotide Sequences**
5. **Probe synthesis**
6. **Protein expression and Purification**
7. **TGT labeling reactions**
8. **Cell culture of U2OS cells**
9. **Northern blotting**
10. **Fluorescence in situ hybridization (FISH)**
11. **Supplementary Figures**
12. **MS of PreQ1 probes**
13. **MS of oligo modifications with various probes**
14. **References**

1. **General Methods, Instrument Details and Materials**

All reagents used for buffers and coupling reactions were purchased from Sigma-Aldrich. Cy5-NHS was purchased from Broadpharm, silicon rhodamine-NHS was purchased from Spirochrome, tetramethylrhodamine-NHS was purchased from Lumiprobe, and biotin-NHS, AlexaFluor647-NHS, and AlexaFluor488-NHS were purchased from Thermo Fisher. All oligonucleotides were purchased from Integrated DNA Technologies. Urea PAGE gels were prepared using the 19:1 sequagel system from National Diagnostics, separated using the Mini-Protean system from Biorad, and stained with GelRed from Biotium. All polyacrylamide gels were imaged on a Bio-Rad ChemiDoc-MP gel imager. Gels images were analyzed using ImageJ software from the NIH. U2OS cells were purchased from ATCC and cultured according to supplier recommendations. Fluorescence in situ hybridization (FISH) probe sets were either purchased from Stellaris or designed using the free online tool on the Stellaris website and ordered from IDT. For IR Northern experiments, the RNA was transferred to a positively charged nylon membrane from Invitrogen and crosslinked on a Stratalinker. His purification was conducted using the HisPur Ni-NTA resin from Thermo Fisher and the His-Spin buffers from Zymo. StrepTag purification was carried out using the Strep-Spin kit from Zymo. Protein gels were run using the Mini-Protean gel system and precast gels from Bio-Rad and stained with InstantBlue Coomassie stain from Abcam. All restriction enzymes, bio-reagents, and competent bacterial strains were purchased from New England Biolabs, Promega, or Life Technologies. Absorbance measurements were obtained with a Thermo Scientific Nano-Drop 2000c UV-Vis spectrophotometer. High resolution mass spectrometry (HRMS) of the small molecule probes and mass spectrometry (MS) of the oligonucleotide samples were collected by the UCSD Department of Chemistry and Biochemistry Molecular Mass Spectroscopy Facility on an Agilent 6230 time-of-flight mass spectrometer (TOFMS) with JetStream electrospray ionization source (ESI).

### 2. **Buffers**

His lysis and wash buffer: 50 mM sodium phosphate buffer pH 7.7, 300 mM sodium chloride, 50 mM imidazole, 0.03 % Triton X-100. (Zymo P2003-4; supplemented with 400 µM PMSF)

His elution buffer: 50 mM sodium phosphate buffer pH 7.7, 300 mM sodium chloride, 250 mM imidazole. (Zymo P2003-5; supplemented with 400 µM PMSF)

TGT storage buffer: 25 mM HEPES, pH 7.5, 100 mM NaCl, 100 µM PMSF.

TGT reaction buffer 1X: 100 mM HEPES, pH 7.3, 5 mM DTT, and 20 mM MgCl2

Northern hybridization buffer: NorthernMax Prehybridization/Hybridization Buffer (Invitrogen AM8677)

Northern wash buffer: Northern Max Low Stringency wash buffer (Invitrogen AM8673)

FISH buffer A: 1:7:2 deionized formamide:water:Stellaris Buffer A (Biosearch Technologies SMF-WA1-60)

FISH buffer B: Stellaris Buffer B (Biosearch Technologies SMF-WB1-20)

FISH hybridization buffer: 9:1 Stellaris Hybridization Buffer:deionized formamide (Biosearch Technologies SMF-HB1-10)

In vitro transcription buffer: 40 mM Tris pH 7.5, 5 mM DTT, 25 mM MgCl2, 2 mM spermidine

### 3. **Plasmid Design and Sequences**

**Plasmid Design:** A second purification tag was added to the TGT-His construct previously reported[2] (Addgene 138201) using the online NEBase changer tool and Q5 site directed mutagenesis kit (NEB E0554S). Two constructs with either an N- or C-terminal Strep purification tag were assembled and used interchangeably. Purification tag sequences are bolded and underlined below. The constructs were transformed into high efficiency 5-alpha competent *E. coli* cells (NEB C2987H), plated, and grown overnight on an LB agar plate containing Kanamycin. The next day, individual colonies were picked, expanded in a 10 mL culture, mini prepped (Biomiga PD1213), and the resulting plasmids sequenced by Eton Biosciences.

**TGT-Strep-His Coding Sequence:** ATGAAATTTGAACTGGACACCACCGACGGTCGCGCACGCCG
TGGCCGCCTGGTCTTTGATCGTGGCGTAGTGGAAACGCCTTGTTTTATGCCTGTTGGCACCTACGGCACC
GTAAAAGGGATGACGCCGGAAGAAGTTGAAGCCACTGGCGCGCAAATTATCCTCGGCAACACCTTCCAC
CTGTGGCTGCGCCCGGGCCAGGAAATCATGAAACTGCACGGCGATCTGCACGATTTTATGCAGTGGAA
GGGGCCGATCCTCACCGACTCCGGCGGCTTCCAGGTCTTCAGCCTTGGCGATATTCGTAAAATCACCGA
ACAGGGCGTGCACTTCCGTAACCCGATCAACGGCGATCCGATTTTCCTCGATCCTGAAAAATCAATGGA
GATTCAGTACGATCTTGGTTCGGATATCGTCATGATCTTTGATGAGTGTACGCCGTATCCTGCTGACTGG
GATTACGCAAAACGCTCCATGGAGATGTCTCTGCGTTGGGCGAAGCGTAGCCGTGAGCGTTTTGACAGT
CTCGGAAACAAAAATGCGCTGTTTGGTATCATCCAGGGCAGCGTTTACGAAGATTTACGTGATATTTCTG
TTAAAGGTCTGGTAGATATCGGTTTTGATGGCTACGCTGTCGGCGGTCTGGCTGTGGGTGAGCCGAAAG
CAGATATGCACCGCATTCTGGAGCATGTATGCCCGCAAATTCCGGCAGACAAACCGCGTTACCTGATGG
GCGTTGGTAAACCAGAAGACCTGGTTGAAGGCGTACGTCGTGGTATCGATATGTTTGACTGCGTAATGC
CAACCCGCAACGCCCGAAATGGTCATTTGTTCGTGACCGATGGCGTGGTGAAAATCCGCAATGCGAAGT
ATAAGAGCGATACTGGCCCACTCGATCCTGAGTGTGATTGCTACACCTGTCGCAATTATTCACGCGCTTA
CTTGCATCATCTTGACCGTTGCAACGAAATATTAGGCGCGCGACTCAACACCATTCATAACCTTCGTTACT
ACCAGCGTTTGATGGCGGGTTTACGCAAGGCTATTGAAGAGGGTAAATTAGAGAGCTTCGTAACTGATT
TTTACCAGCGTCAGGGGCGAGAAGTACCACCTTTGAACGTTGATGGC**TGGAGCCACCCGCAGTTCGAA**
**AAA**CTCGAG**CACCATCACCACCATCAC**TAA

**Strep-TGT-His Coding Sequence:** ATG**TGGAGCCATCCGCAGTTTGAAAAA**GGCAAATTTGAACT
GGACACCACCGACGGTCGCGCACGCCGTGGCCGCCTGGTCTTTGATCGTGGCGTAGTGGAAACGCCTTG
TTTTATGCCTGTTGGCACCTACGGCACCGTAAAAGGGATGACGCCGGAAGAAGTTGAAGCCACTGGCGC
GCAAATTATCCTCGGCAACACCTTCCACCTGTGGCTGCGCCCGGGCCAGGAAATCATGAAACTGCACGG
CGATCTGCACGATTTTATGCAGTGGAAGGGGCCGATCCTCACCGACTCCGGCGGCTTCCAGGTCTTCAG
CCTTGGCGATATTCGTAAAATCACCGAACAGGGCGTGCACTTCCGTAACCCGATCAACGGCGATCCGAT
TTTCCTCGATCCTGAAAAATCAATGGAGATTCAGTACGATCTTGGTTCGGATATCGTCATGATCTTTGATG
AGTGTACGCCGTATCCTGCTGACTGGGATTACGCAAAACGCTCCATGGAGATGTCTCTGCGTTGGGCGA
AGCGTAGCCGTGAGCGTTTTGACAGTCTCGGAAACAAAAATGCGCTGTTTGGTATCATCCAGGGCAGCG
TTTACGAAGATTTACGTGATATTTCTGTTAAAGGTCTGGTAGATATCGGTTTTGATGGCTACGCTGTCGG
CGGTCTGGCTGTGGGTGAGCCGAAAGCAGATATGCACCGCATTCTGGAGCATGTATGCCCGCAAATTCC
GGCAGACAAACCGCGTTACCTGATGGGCGTTGGTAAACCAGAAGACCTGGTTGAAGGCGTACGTCGTG
GTATCGATATGTTTGACTGCGTAATGCCAACCCGCAACGCCCGAAATGGTCATTTGTTCGTGACCGATGG
CGTGGTGAAAATCCGCAATGCGAAGTATAAGAGCGATACTGGCCCACTCGATCCTGAGTGTGATTGCTA
CACCTGTCGCAATTATTCACGCGCTTACTTGCATCATCTTGACCGTTGCAACGAAATATTAGGCGCGCGA
CTCAACACCATTCATAACCTTCGTTACTACCAGCGTTTGATGGCGGGTTTACGCAAGGCTATTGAAGAGG
GTAAATTAGAGAGCTTCGTAACTGATTTTTACCAGCGTCAGGGGCGAGAAGTACCACCTTTGAACGTTG
AT**CACCATCACCACCATCAC**TAA

4. **Oligonucleotides**

#### Key hairpins

**Table S1.** Key hairpins referenced in this work. RNA hairpins are shown in blue. DNA hairpins are shown in black.

| Design | Hairpin | Sequence |
|---|---|---|
| original hairpins | ECY-A1 | GCAGACUGUAAAUCUGC |
| | ECYMH | GGGAGCAGACUGUAAAUCUGCUCCC |
| mini helix mutants | ECYMH-rTGrT | GGGAGCAGACrTGrTAAArTCrTGCrTCCC |
| | dECYMH | GGGAGCAGACTGTAAATCTGCTCCC |
| | dECYMH-dUGdU | GGGAGCAGACdUGdUAAATCTGCTCCC |
| | dECYMH-dUGT | GGGAGCAGACdUGTAAATCTGCTCCC |
| | dECYMH-TGdU | GGGAGCAGACTGdUAAATCTGCTCCC |
| tRNA analogs | dECY-0 (dECY-A1) | GCAGACTGTAAATCTGC |
| | dECH-0 | CTGGATTGTGATTCCAG |
| tRNA chimera | dECH-dECD | CTGGACTGTCACTCCAG |
| top dECH-9 loop mutants | dECH-10 | CTGGATTGTCCTTCCAG |
| | dECH-18 | CTGGACTGTCCCTCCAG |
| | dECH-24 | CTGGACTGTTCCTCCAG |
| | dECH-26 | CTGGATTGTCCCTCCAG |
| | dECH-32 | CTGGATTGTTCCTCCAG |
| | dECH-34 | CTGGACTGTCCTTCCAG |
| ΔC mutant | dECH-10 ΔC | CTGGATTCTCCTTCCAG |
| top stem sequence mutants dECH-10 loop | 162_ala2_dECH-10 | CCTGCTTGTCCTGCAGG |
| | 164_arg2_dECH-10 | CTCGGTTGTCCTCCGAG |
| | 165_arg3_dECH-10 | CTGCCTTGTCCTGGCAG |
| | 170_gln1_dECH-10 | CCGGATTGTCCTTCCGG |
| | 171_gln2_dECH-10 | CCGGTTTGTCCTACCGG |
| | 172_glu1_dECH-10 | CCGCCTTGTCCTGGCGG |
| | 174_gly2_dECH-10 | TCAGCTTGTCCTGCTGA |
| | 175_gly3_dECH-10 | CGACCTTGTCCTGGTCG |
| | 176_his1_dECH-10 | CTGGATTGTCCTTCCAG |
| | 177_ile1_dECH-10 | CACCCTTGTCCTGGGTG |
| | 180_leu3_dECH-10 | CTACCTTGTCCTGGTAG |
| | 194_ser3_dECH-10 | CTCCCTTGTCCTGGGAG |
| | 197_thr2_dECH-10 | TCGCATTGTCCTTGCGA |
| | 198_thr3_dECH-10 | CGCATTTGTCCTATGCG |
| | 203_val1_dECH-10 | CCACCTTGTCCTGGTGG |
| | 204_val2_dECH-10 | CCTCCTTGTCCTGGAGG |

### Analog Hairpins

**Table S2.** Analog hairpins based on known constructs. RNA hairpins are shown in blue. DNA hairpins are shown in black. Key hairpins are also included in this table.

| Design | Hairpin | Sequence |
|---|---|---|
| original hairpins | rECY-A1 | GCAGACUGUAAAUCUGC |
| | rECYMH | GGGAGCAGACUGUAAAUCUGCUCCC |
| mini helix mutants | rECYMH-rTGrT | GGGAGCAGACrTGrTAAArTCrTGCrTCCC |
| | dECYMH | GGGAGCAGACTGTAAATCTGCTCCC |
| | dECYMH-dUGdU | GGGAGCAGACdUGdUAAATCTGCTCCC |
| | dECYMH-dUGT | GGGAGCAGACdUGTAAATCTGCTCCC |
| | dECYMH-TGdU | GGGAGCAGACTGdUAAATCTGCTCCC |
| tRNA analogs | dECY-0 (dECY-A1) | GCAGACTGTAAATCTGC |
| | dECH-0 | CTGGATTGTGATTCCAG |
| | dECD-0 | CCTGCCTGTCACGCAGG |
| | dECN-0 | GCGGACTGTTAATCCGT |
| tRNA chimeras | dECD-dECH | CCTGCTTGTGATGCAGG |
| | dECD-dECY | CCTGCCTGTAAAGCAGG |
| | dECD-dECN | CCTGCCTGTTAAGCAGG |
| | dECN-dECH | GCGGATTGTGATTCCGT |
| | dECN-dECY | GCGGACTGTAAATCCGT |
| | dECN-dECD | GCGGACTGTCACTCCGT |
| | dECY-dECH | GCAGATTGTGATTCTGC |
| | dECY-dECN | GCAGACTGTTAATCTGC |
| | dECY-dECD | GCAGACTGTCACTCTGC |
| | dECH-dECY | CTGGACTGTAAATCCAG |
| | dECH-dECN | CTGGACTGTTAATCCAG |
| | dECH-dECD | CTGGACTGTCACTCCAG |

### Mutant Hairpins

**Table S3.** Loop and stem mutant hairpins based on anticodon arms of tRNAs. Redundant hairpins are group by color. Key hairpins are also included in these tables.

| | Hairpin | Sequence |
|---|---|---|
| dECH mutants | dECH-1 | CTCGATTGTGATTCGAG |
| | dECH-2 | CTAGATTGTGATTCTAG |
| | **dECH-3** | **CTGGATTGTTATTCCAG** |
| | dECH-4 | CTGCATTGTGATTGCAG |
| | dECH-5 | CTGAATTGTGATTTCAG |
| | dECH-6 | GTGGATTGTGATTCCAC |
| | dECH-7 | GTGGATTGTGATTCCAT |
| | dECH-8 | ATGGATTGTGATTCCAT |
| | dECH-9 | CTGGATTGTCATTCCAG |
| | dECH-10 | CTGGATTGTCCTTCCAG |
| | dECH-11 | CTGGATTGTTTTTCCAG |
| | **dECH-12** | **CTGGATTGTTATTCCAG** |
| | dECH-13 | CTGGATTGTCTTTCCAG |
| | dECH-14 | CTGGATTGTCGTTCCAG |
| | dECH-15 | CTGGATTGTTGTTCCAG |
| | dECH-16 | CTGGATTGTTCTTCCAG |
| | dECH-17 | CTGGACTGTCACTCCAG |
| | dECH-18 | CTGGACTGTCCCTCCAG |
| | dECH-19 | CTGGACTGTTTCTCCAG |
| | dECH-20 | CTGGACTGTTACTCCAG |
| | dECH-21 | CTGGACTGTCTCTCCAG |
| | dECH-22 | CTGGACTGTCGCTCCAG |
| | dECH-23 | CTGGACTGTTGCTCCAG |
| | dECH-24 | CTGGACTGTTCCTCCAG |
| | dECH-25 | CTGGATTGTCACTCCAG |
| | dECH-26 | CTGGATTGTCCCTCCAG |
| | dECH-27 | CTGGATTGTTTCTCCAG |
| | dECH-28 | CTGGATTGTTACTCCAG |
| | dECH-29 | CTGGATTGTCTCTCCAG |
| | dECH-30 | CTGGATTGTCGCTCCAG |
| | dECH-31 | CTGGATTGTTGCTCCAG |
| | dECH-32 | CTGGATTGTTCCTCCAG |
| | dECH-33 | CTGGACTGTCATTCCAG |
| | dECH-34 | CTGGACTGTCCTTCCAG |
| | dECH-35 | CTGGACTGTTTTTCCAG |
| | dECH-36 | CTGGACTGTTATTCCAG |
| | dECH-37 | CTGGACTGTCTTTCCAG |
| | dECH-38 | CTGGACTGTCGTTCCAG |
| | dECH-39 | CTGGACTGTTGTTCCAG |
| | dECH-40 | CTGGACTGTTCTTCCAG |

| | Hairpin | Sequence |
|---|---|---|
| dECH-10 loop/tRNA stem mutants | 161_ala1_dECH-10 | CTTGCTTGTCCTGCAAG |
| | **162_ala2_dECH-10** | **CCTGCTTGTCCTGCAGG** |
| | 163_arg1_dECH-10 | AGCCCTTGTCCTGGGCT |
| | 164_arg2_dECH-10 | CTCGGTTGTCCTCCGAG |
| | 165_arg3_dECH-10 | CTGCCTTGTCCTGGCAG |
| | 166_arg4_dECH-10 | ACGACTTGTCCTGTCGT |
| | **167_asn1_dECH-10** | **GCGGATTGTCCTTCCGT** |
| | **168_asp1_dECH-10** | **CCTGCTTGTCCTGCAGG** |
| | **169_cys1_dECH-10** | **GCGGATTGTCCTTCCGT** |
| | 170_gln1_dECH-10 | CCGGATTGTCCTTCCGG |
| | **171_gln2_dECH-10** | **CCGGTTTGTCCTACCGG** |
| | 172_glu1_dECH-10 | CCGCCTTGTCCTGGCGG |
| | 173_gly1_dECH-10 | AGAGCTTGTCCTGCTCT |
| | 174_gly2_dECH-10 | TCAGCTTGTCCTGCTGA |
| | 175_gly3_dECH-10 | CGACCTTGTCCTGGTCG |
| | 176_his1_dECH-10 | CTGGATTGTCCTTCCAG |
| | **177_ile1_dECH-10** | **CACCCTTGTCCTGGGTG** |
| | 178_leu1_dECH-10 | GTTGATTGTCCTTCAAC |
| | 179_leu2_dECH-10 | CTAGCTTGTCCTGTTAG |
| | 180_leu3_dECH-10 | CTACCTTGTCCTGGTAG |
| | 181_leu4_dECH-10 | CCAGATTGTCCTTCTGG |
| | 182_leu5_dECH-10 | AGGGATTGTCCTTCCCT |
| | 183_lys1_dECH-10 | GTTGATTGTCCTTCAAT |
| | **184_met1_dECH-10** | **GGCGATTGTCCTTCGCT** |
| | 185_met2_dECH-10 | CATCATTGTCCTTGATG |
| | **186_met3_dECH-10** | **GGCGATTGTCCTTCGCT** |
| | 187_met4_dECH-10 | TCGGGTTGTCCTCCCGA |
| | 188_phe1_dECH-10 | GGGGATTGTCCTTCCCC |
| | 189_pro1_dECH-10 | ACTGGTTGTCCTCCAGT |
| | 190_pro2_dECH-10 | CCGTCTTGTCCTGTCGG |
| | 191_pro3_dECH-10 | CTTCGTTGTCCTCGAGG |
| | **192_ser1_dECH-10** | **CCGGTTTGTCCTACCGG** |
| | **193_ser2_dECH-10** | **CCGGTTTGTCCTACCGG** |
| | 194_ser3_dECH-10 | CTCCCTTGTCCTGGGAG |
| | 195_ser4_dECH-10 | CACGCTTGTCCTGTGTG |
| | **196_thr1_dECH-10** | **CACCCTTGTCCTGGGTG** |
| | 197_thr2_dECH-10 | TCGCATTGTCCTTGCGA |
| | 198_thr3_dECH-10 | CGCATTTGTCCTATGCG |
| | 199_thr4_dECH-10 | ACTGATTGTCCTTCAGT |
| | **200_thr5_dECH-10** | **CACCCTTGTCCTGGGTG** |
| | **201_trp1_dECH-10** | **CCGGTTTGTCCTACCGG** |
| | 202_tyr1_dECH-10 | GCAGATTGTCCTTCTGC |
| | 203_val1_dECH-10 | CCACCTTGTCCTGGTGG |
| | 204_val2_dECH-10 | CCTCCTTGTCCTGGAGG |

### Oligo Extension Hairpins

**Table S4.** Labeling of oligos containing one or two recognition hairpins. DNA-TAG recognition hairpins are underlined.

| Oligo Extension Hairpins | | |
|---|---|---|
| **Design** | **Hairpin** | **Sequence** |
| single hairpin | His M10 T2 int | ACCATCATTTTCATATCCTCCActggaTTGTCCTtccagACCACCATCATTTGCAATGA |
| | His M10 T2 3' | ACCATCATTTTCATATCCTCCAACCACCATCATTTGCAATGActggaTTGTCCTtccag |
| | His M10 T2 5' | ctggaTTGTCCTtccagACCATCATTTTCATATCCTCCAACCACCATCATTTGCAATGA |
| double hairpin | 5/3_176/177_**GG** | ctggaTT**G**TCCTtccagCATCATTTTCATATCCTCCAcacccTT**G**TCCTgggtg |
| | 5/3_176/177_**GC** | ctggaTT**G**TCCTtccagCATCATTTTCATATCCTCCAcacccTT**C**TCCTgggtg |
| | 5/3_176/177_**CG** | ctggaTT**C**TCCTtccagCATCATTTTCATATCCTCCAcacccTT**G**TCCTgggtg |
| | 5/5_176/177_**GG** | ctggaTT**G**TCCTtccagTTTcacccTT**G**TCCTgggtgCATCATTTTCATATCCTCCA |
| | 5/5_176/177_**GC** | ctggaTT**G**TCCTtccagTTTcacccTT**C**TCCTgggtgCATCATTTTCATATCCTCCA |
| | 5/5_176/177_**CG** | ctggaTT**C**TCCTtccagTTTcacccTT**G**TCCTgggtgCATCATTTTCATATCCTCCA |
| | 3/3_176/177_**GG** | CATCATTTTCATATCCTCCActggaTT**G**TCCTtccagTTTcacccTT**G**TCCTgggtg |
| | 3/3_176/177_**GC** | CATCATTTTCATATCCTCCActggaTT**G**TCCTtccagTTTcacccTT**C**TCCTgggtg |
| | 3/3_176/177_**CG** | CATCATTTTCATATCCTCCActggaTT**C**TCCTtccagTTTcacccTT**G**TCCTgggtg |

**In vitro transcription (IVT) of ECYMH-rTGrT.** Forward and reverse template constructs were ordered from IDT and annealed (Table S5). The transcription was carried out with the following conditions using T7 RNA polymerase prepared as previously described.[1] 1X T7 RNAP Buffer, 5 mM MeUTP, 5 mM ATP, 5 mM CTP, 9 mM GTP, 5 mM annealed template, 0.004 U/µL PPTase, 0.05% Triton X, and 0.15 µg/µL T7 RNAP and incubated at 4˚C for 4 hours. Subsequently, 1.2 µL 100 mM $CaCl_2$ and 1 µL Turbo DNAse (Thermo Fisher AM2238) were added to a 100 µL IVT reaction and incubated at 37˚C for 20 minutes. The RNA product was then purified by ethanol precipitation and the transcript analyzed by urea PAGE.

**Table S5.** IVT template for rECYMH-rTGrT. The T7 promoter sequence is underlined, and the hairpin sequence is in bold.

| **Construct** | **Sequence** |
|---|---|
| **rECYMH-rTGrT FWD** | GAAATTAATACGACTCACTATA**GGGAGCAGACTGTAAATCTGCTCCC** |
| **rECYMH-rTGrT REV** | **GGGAGCAGATTTACAGTCTGCTCCC**TATAGTGAGTCGTATTAATTTC |

5. **Probe Synthesis**

All probes were synthesized as described previously by removing a tert-butyloxycarbonyl (boc) protecting group from a preQ1 derivative and coupling it to commercially available NHS functionalized reporters.[1,2] Briefly, the boc protected starting material was deprotected in 10% TFA/DCM for one hour. The TFA was neutralized 3 times with 10% TEA in DCM, rotoevaporating the solution to near dryness between each wash. The crude oil product was taken to the next step where it was dissolved in dry DMF followed by the dropwise addition of TEA. Each NHS probe was dissolved in dry DMF and added to the reaction dropwise with stirring. The reaction was allowed to proceed for 2 hours at room temperature, the DMF evaporated, and the product purified by HPLC. The final products were confirmed by LCMS.

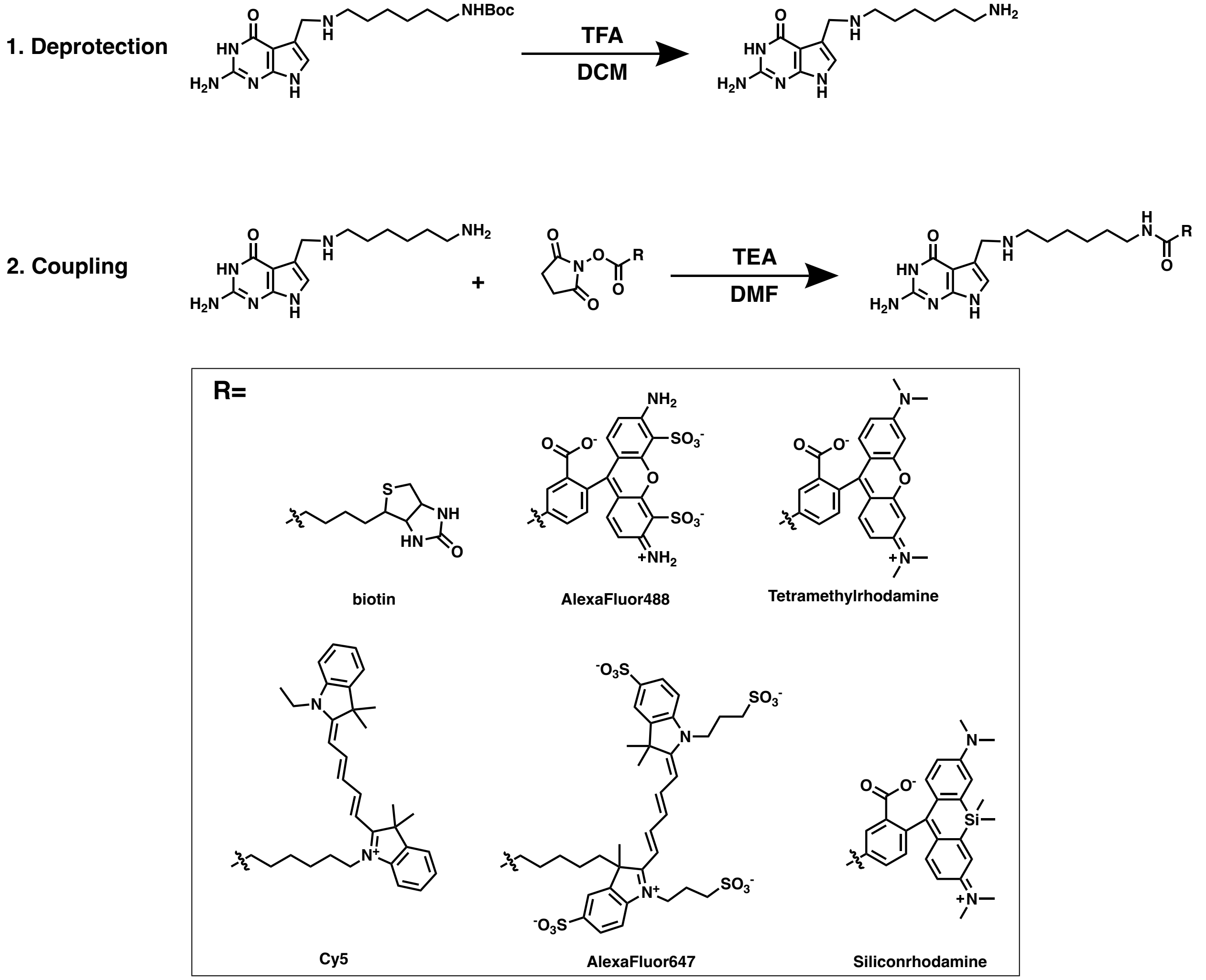


**Scheme S1.** Synthesis of preQ1 probes with commercially purchased preQ1-$C_6$NHBoc.

6. **Protein Expression and Purification**

**Expression of TGT** *E. coli* TGT was expressed as previously described from the plasmid described in section 3.[2] Briefly, the plasmid carrying TGT was transformed into BL21(DE3) Competent *E. coli* cells (NEB C2527) and a starter culture was grown overnight. Approximately 5 mL of starter culture was transferred into 200 mL of kanamycin containing LB and allowed to grow to an optical density of ~0.6. At this point, expression was induced with by adding IPTG to a final concentration of 1 mM and shaken at 37˚C for 4 hours. The bacteria were pelleted by centrifugation (6,000xg for 20 minutes at 4˚C). Cell pellets were either resuspended and purified immediately or stored at -20˚C if desired.

**Purification of TGT** A dual purification (His and streptactin) protocol was used to increase purity of the enzyme. 10µL aliquots were taken at each step for analysis via 4-20% SDS PAGE.

***His purification*** was carried out using HisPur™ Ni-NTA Resin spin columns (Thermo Fisher 88224) and His purification buffers (Zymo P2003-4; P2003-5). The cell pellet was resuspended in 2 mL total of lysis buffer and sonicated on ice using the following settings: 4 cycles (30 sec on / 2 min off); output control = 4; duty cycle% = 30-40. The lysate was centrifuged at 10,000xg for 30 min at 4˚C to remove cellular debris. In tandem, the resin from the spin column was equilibrated 2 times with 400 µL of lysis buffer. The supernatant of the centrifuged cell lysate was transferred to the capped spin column and incubated for at least one hour at 4˚C with end over end mixing. The resin was then washed and the protein eluted according to the HisPur manual. The lysis and wash buffer are the same as it was noted that a higher imidazole concentration during binding reduced nonspecific carry over into the final elutions. The elution was taken directly to streptactin purification.

***Streptactin purification*** was carried out using the Strep-Spin Protein Miniprep Kit (Zymo P2004). It should be noted that the protein sample incubation time was increased to 30-60 minutes at 4˚C to increase yield. After elution from the streptactin column, the enzyme prep was dialyzed against TGT storage buffer to remove biotin and concentrated using a 30 kDa cut off spin filter (Millipore UFC503096). Enzyme concentration was determined using a BCA assay kit (Thermo Fisher 23227). The final protein was aliquoted in 20 µL fractions and stored at -80˚C.

7. **TGT labeling and analysis**

**TGT labeling reaction:** TGT modification of DNA and RNA substrates was carried out at 37˚C for 4 hours in a thermocycler with a heated lid[1,2] using the following reaction conditions: 10 µM enzyme, 5 µM nucleic acid substrate, and 50 µM of preQ1-X (various preQ1 based probes), 1X TGT reaction buffer, and 1-unit RNAsin from Promega for RNA substrates. Components were mixed together, gently vortexed, briefly spun down, and incubated on a thermocycler. For labeling reactions of constructs containing 2 hairpins, the small molecule substrate concentration was doubled. For analysis purposes, reactions were not purified before PAGE, which did not affect the gel shift.

**Labeled oligo purification:** For the experiments where it was necessary to purify the labeled construct, an oligo clean and concentrator kit (Zymo D4060) was used, and the product eluted in pure water.

**Urea PAGE:** Urea PAGE was the primary method of analysis to determine TGT activity toward specific substrates. For all gels, we used the 19:1 Acrylamide:Bisacrylamide SequaGel® UreaGel System (National Diagnostics EC-833) and the Mini-PROTEAN Tetra Vertical Electrophoresis Cell from Bio- Rad. Briefly, 1.5 mm plates assembled on the Bio-Rad gel casting stand, 10 mL of the desired percentage gel was freshly prepared in a 15 mL centrifuge tube, 14 µL of TEMED was added and gently mixed by inverting the tube, followed by 80 µL of 10% ammonium per sulfate. After briefly mixing, the solution is poured between the plates, a comb inserted, and allowed to solidify (~20-30 min). All oligo samples were prepared for gel electrophoresis using 2X RNA loading dye (NEB B0363S). For all constructs, 150-200 ng of the oligonucleotide was loaded onto the gel for visualization. The solutions were denatured at 98˚C for 5 minutes on a thermocycler. The electrophoresis cassette was assembled, and the chamber filled with 1X TBE. After thorough washing of the wells to remove leached urea, the samples were loaded into the wells and the gel was run for 10 min at 100V then 90 min at 200V. Afterward, the gel was stained with gel red in TBE (1:1000 dilution; Biotium 41011) for ~5 min and imaged on a Bio-Rad ChemiDoc gel imager system.

**Gel Shift Analysis:** Images were exported for analysis form the ChemiDoc software and analyzed qualitatively by eye. Successful labeling is indicated by an upward shift in the product lane due to an increase in molecular weight of the labeled species. In the cases where a quantitative measurement was desired, the bands were analyzed via densitometry using ImageJ. Briefly, the percent signal of each band compared to the total in the control lane was calculated.

8. **Cell Culture of U2OS Cells**

For Northern and FISH experiments, we used U2OS cells, a human osteosarcoma derived line. The cells were ordered from ATCC (HTB-96), thawed, expanded, and stocks stored at -80˚C. Cells were cultured according to ATCC guidelines in T25 flasks. Complete growth medium containing DMEM (Gibco 11995065), 10% FBS, and 1% pen/strep (Gibco 15070063) was used to maintain and culture the cells. The cells were passaged ~3 times per week at a 1:6 sub cultivation ratio when they reached ~80% confluency. To passage the cells, they were washed 1X with HBSS (Gibco 14025076) and treated with TrypLE Express (Gibco 12604013) at 37˚C until fully lifted. The trypsin was quenched by adding an equal volume of complete medium and the necessary volume of cells taken into a new flask. For experiments, cells were seeded in a black glass bottom 24 well plate at a 1:6 ratio calculated to scale with the area of the plate.

### 9. **Northern blot**

DNA was extracted from U2OS cells and the U6 sequence amplified with a primer set that includes the T7 promoter sequence. This amplified product was purified and used as a template for in vitro transcription. The IVT RNA was purified by ethanol precipitation and analyzed via urea PAGE. Total RNA was extracted from U2OS cells. The U6 PCR amplified template, IVT U6 RNA, and total extracted U2OS RNA were analyzed via gel electrophoresis.

**Table S6.** Relevant sequences for U6 RNA northern blot. The target and antisense sequences are underlined and the DNA-TAG recognition hairpin is in bold.

| Construct | Sequence |
|---|---|
| **U6 RNA** | GUGCUCGCUUCGGCAGCACAUAUACUAAAAUUGGAA<u>CGAUACAGAGAAGAUUAGCAUG</u><br><u>GCCCCUGC</u>GCAAGGAUGACACGCAAAUUCGUGAAGCGUUCCAUAUUUU |
| **5′ dECH-10 U6 probe** | **CTGGATTGTCCTTCCAG**TTT<u>GCAGGGGCCATGCTAATCTTCTCTGTATCG</u> |
| **T7 U6 amp fwd** | ATACGACTCACTATAGGTGCTCGCTTCGGCAG |
| **T7 U6 amp rev** | AAAATATGGAACGCTTCACGAATTTGCGTGTC |

**U6 RNA Template Preparation** Genomic DNA was extracted from U2OS cells using the Quick-DNA Miniprep Plus Kit (Zymo D4068). The U6 RNA IVT template was PCR amplified from ~10 ng of the isolated genomic DNA using Q5® Hot Start High-Fidelity 2X Master Mix (NEB M0494). The PCR product was verified by gel electrophoresis, purified using a DNA clean and concentrator kit (Zymo D4029), and used for the IVT of U6 RNA.

**IVT (In vitro Transcription)** The U6 RNA transcript was *in vitro* transcribed with the following conditions using T7 RNA polymerase prepared as previously described:[1] 1X T7 RNAP Buffer, 5 mM UTP, 5 mM ATP, 5 mM CTP, 9 mM GTP, 5 mM Annealed template, 0.004 U/µL PPTase, 0.05% Triton X, and 0.15 µg/µL T7 RNAP and incubated at 4˚C for 4 hours. Subsequently, 1.2 µL 100 mM CaCl2 and 1 µL Turbo DNAse (Thermo Fisher AM2238) were added to a 100 µL IVT reaction, incubated at 37˚C for 20 minutes. The RNA product was then purified by ethanol precipitation and the transcript analyzed by urea PAGE.

**RNA Extraction** Total RNA was extracted from U2OS cells using the Direct-zol RNA Miniprep Kit (Zymo R2053) and analyzed by urea PAGE.

**Northern Blot** Dilution series of in vitro transcribed U6 RNA or U2OS total RNA were run on a 15% urea PAGE gel as described above. The RNA was transferred to a positively charged nylon membrane (Invitrogen AM10100) using the Mini Trans-Blot cell system from Bio-Rad. The transfer was done at room temperature and run at 200 mA for 30 min in 0.5X TBE. The membrane was washed in 10X SSC buffer for 10 minutes with rocking and crosslinked twice on a Stratalinker using the auto setting. The membrane was then incubated with the 5′ SiR labeled U6 antisense probe in Northern Hybridization Buffer (10 picomolar; 2hr 37˚C), washed 2X for 5 minutes each with Northern Max Low Stringency wash buffer, and imaged on a fluorescent gel scanner. The signal was quantified via densitometry using ImageJ and analyzed for linear fit.

10. **RNA Fluorescence in situ Hybridization (FISH)**

U2OS cells were seeded in glass bottom black 24 well plates at a ratio of 1:6 and allowed to grow to ~80% confluency. Cells were washed with HBSS and fixed in 4% para formaldehyde (PFA) for 10 minutes at room temperature. PFA was aspirated, the cells were washed twice with HBSS, and permeabilized with 70% ethanol overnight at 4˚C. The ethanol solution was removed, and the cells washed 2X with HBSS. FISH buffer A was added and let sit for 5 minutes. Fluorescent probe sets were prepared in hybridization buffer with a final concentration of 125 nM and added to the wells. Parafilm was placed over the wells and around the lid of the plate to minimize evaporation and the plate incubated at 37˚C overnight in the dark. The next morning, the probe solution was aspirated, and the cells washed in FISH buffer A for 30 min at 37˚C in the dark. FISH buffer A was then aspirated from the cells, a solution of 5 ng/mL DAPI in FISH buffer A was added, and the plate was placed at 37˚C for 30 minutes. The DAPI solution was aspirated, and the cells washed in FISH buffer B for 5 minutes. FISH buffer B was aspirated and Vectashield (Vector Laboratories H-1000-10) added to the wells. The cells were imaged on a confocal microscope. All images were window leveled and analyzed visually using ImageJ. For wells with multiple probe sets, equal amounts of each set were added to a final total concentration of 125 nM.

**Table S7.** MALAT 1 single label probe set.

| Probe | Sequence |
|---|---|
| **MALAT1_1_176/177_3'3'_GC** | GCCCTTCTATTGGTATTAATTTTCTGGATTGTCCTTCCAGTTTCACCCTTCTCCTGGGTG |
| **MALAT1_2_176/177_3'3'_GC** | CTCCCAATTAATCTTTCCATTTTCTGGATTGTCCTTCCAGTTTCACCCTTCTCCTGGGTG |
| **MALAT1_3_176/177_3'3'_GC** | TCTCCAAATTGTTTCATCCTTTTCTGGATTGTCCTTCCAGTTTCACCCTTCTCCTGGGTG |
| **MALAT1_4_176/177_3'3'_GC** | TTAGCTTTTTGTTTCCTAGCTTTCTGGATTGTCCTTCCAGTTTCACCCTTCTCCTGGGTG |
| **MALAT1_5_176/177_3'3'_GC** | ATCTTCTCAAGCTTTACCTTTTTCTGGATTGTCCTTCCAGTTTCACCCTTCTCCTGGGTG |
| **MALAT1_6_176/177_3'3'_GC** | ACTATATTTAAGGCCTTCCATTTCTGGATTGTCCTTCCAGTTTCACCCTTCTCCTGGGTG |
| **MALAT1_7_176/177_3'3'_GC** | TACTTCCGTTACGAAAGTCCTTTCTGGATTGTCCTTCCAGTTTCACCCTTCTCCTGGGTG |
| **MALAT1_8_176/177_3'3'_GC** | CTGGGTCAGCTGTCAATTAATTTCTGGATTGTCCTTCCAGTTTCACCCTTCTCCTGGGTG |
| **MALAT1_9_176/177_3'3'_GC** | TCAGTCCTAGCTTCATCAAATTTCTGGATTGTCCTTCCAGTTTCACCCTTCTCCTGGGTG |
| **MALAT1_10_176/177_3'3'_GC** | AACAACATATTGCCGACCTCTTTCTGGATTGTCCTTCCAGTTTCACCCTTCTCCTGGGTG |
| **MALAT1_11_176/177_3'3'_GC** | AGTCATTTGCCTTTAGGATTTTTCTGGATTGTCCTTCCAGTTTCACCCTTCTCCTGGGTG |
| **MALAT1_12_176/177_3'3'_GC** | AACTGTAAACCTGTGGTGGTTTTCTGGATTGTCCTTCCAGTTTCACCCTTCTCCTGGGTG |
| **MALAT1_13_176/177_3'3'_GC** | CTCAACGTGAGAACTGCTCTTTTCTGGATTGTCCTTCCAGTTTCACCCTTCTCCTGGGTG |
| **MALAT1_14_176/177_3'3'_GC** | CTCTAACCCAGTTTGTCAATTTTCTGGATTGTCCTTCCAGTTTCACCCTTCTCCTGGGTG |
| **MALAT1_15_176/177_3'3'_GC** | CCAAGGATAAAAGCAGCTCCTTTCTGGATTGTCCTTCCAGTTTCACCCTTCTCCTGGGTG |
| **MALAT1_16_176/177_3'3'_GC** | TGAACCAAAGCTGCACTGTGTTTCTGGATTGTCCTTCCAGTTTCACCCTTCTCCTGGGTG |
| **MALAT1_17_176/177_3'3'_GC** | ACTGCCAACTAATTGCCAATTTTCTGGATTGTCCTTCCAGTTTCACCCTTCTCCTGGGTG |
| **MALAT1_18_176/177_3'3'_GC** | CCAGTGGCTCATATTTAACTTTTCTGGATTGTCCTTCCAGTTTCACCCTTCTCCTGGGTG |
| **MALAT1_19_176/177_3'3'_GC** | ACTTTCCTTGCCCAAATTAATTTCTGGATTGTCCTTCCAGTTTCACCCTTCTCCTGGGTG |
| **MALAT1_20_176/177_3'3'_GC** | CCCAATGGAGGTATGACATATTTCTGGATTGTCCTTCCAGTTTCACCCTTCTCCTGGGTG |
| **MALAT1_21_176/177_3'3'_GC** | CATGCAATACTGCAGATGCATTTCTGGATTGTCCTTCCAGTTTCACCCTTCTCCTGGGTG |
| **MALAT1_22_176/177_3'3'_GC** | AGGAGAAAGTGCCATGGTTGTTTCTGGATTGTCCTTCCAGTTTCACCCTTCTCCTGGGTG |
| **MALAT1_23_176/177_3'3'_GC** | TTTCTCAATCCTGAAATCCCTTTCTGGATTGTCCTTCCAGTTTCACCCTTCTCCTGGGTG |
| **MALAT1_24_176/177_3'3'_GC** | TGAAGTGTACTATCCCATCATTTCTGGATTGTCCTTCCAGTTTCACCCTTCTCCTGGGTG |
| **MALAT1_25_176/177_3'3'_GC** | TCCTGATCTGGTCCATTAAATTTCTGGATTGTCCTTCCAGTTTCACCCTTCTCCTGGGTG |
| **MALAT1_26_176/177_3'3'_GC** | TCTTTCCTGCCTTAAAGTTATTTCTGGATTGTCCTTCCAGTTTCACCCTTCTCCTGGGTG |
| **MALAT1_27_176/177_3'3'_GC** | TGTCAATTTATAGACCCCTGTTTCTGGATTGTCCTTCCAGTTTCACCCTTCTCCTGGGTG |
| **MALAT1_28_176/177_3'3'_GC** | AAACATTGCCTACCACTCTATTTCTGGATTGTCCTTCCAGTTTCACCCTTCTCCTGGGTG |
| **MALAT1_29_176/177_3'3'_GC** | CCTGAATGGCTTCATGAAGGTTTCTGGATTGTCCTTCCAGTTTCACCCTTCTCCTGGGTG |
| **MALAT1_30_176/177_3'3'_GC** | TGCATTTACTTGCCAACAGATTTCTGGATTGTCCTTCCAGTTTCACCCTTCTCCTGGGTG |
| **MALAT1_31_176/177_3'3'_GC** | GTCGTTTCACAATGCATTCTTTTCTGGATTGTCCTTCCAGTTTCACCCTTCTCCTGGGTG |
| **MALAT1_32_176/177_3'3'_GC** | CAACACTCAGCCTTTATCACTTTCTGGATTGTCCTTCCAGTTTCACCCTTCTCCTGGGTG |
| **MALAT1_33_176/177_3'3'_GC** | TTTTTTCTTACTGGGTCTGGTTTCTGGATTGTCCTTCCAGTTTCACCCTTCTCCTGGGTG |
| **MALAT1_34_176/177_3'3'_GC** | ACTGGAAGCTCCTTCTATAGTTTCTGGATTGTCCTTCCAGTTTCACCCTTCTCCTGGGTG |
| **MALAT1_35_176/177_3'3'_GC** | TCATTTTGTCCACTGGTGAATTTCTGGATTGTCCTTCCAGTTTCACCCTTCTCCTGGGTG |
| **MALAT1_36_176/177_3'3'_GC** | TTGTCCCATAACTGATCTGATTTCTGGATTGTCCTTCCAGTTTCACCCTTCTCCTGGGTG |
| **MALAT1_37_176/177_3'3'_GC** | AACACAGTTTGCTCACATGCTTTCTGGATTGTCCTTCCAGTTTCACCCTTCTCCTGGGTG |
| **MALAT1_38_176/177_3'3'_GC** | TGACACTTCTCTTGACCTTATTTCTGGATTGTCCTTCCAGTTTCACCCTTCTCCTGGGTG |
| **MALAT1_39_176/177_3'3'_GC** | CACTCCAGAAAGAGGGAGTTTTTCTGGATTGTCCTTCCAGTTTCACCCTTCTCCTGGGTG |
| **MALAT1_40_176/177_3'3'_GC** | AGATCAAAAGGCACGGGGTGTTTCTGGATTGTCCTTCCAGTTTCACCCTTCTCCTGGGTG |
| **MALAT1_41_176/177_3'3'_GC** | ACAAGGATCCAAGCTACTGGTTTCTGGATTGTCCTTCCAGTTTCACCCTTCTCCTGGGTG |
| **MALAT1_42_176/177_3'3'_GC** | CATCGTTACCTTGAAACCGATTTCTGGATTGTCCTTCCAGTTTCACCCTTCTCCTGGGTG |
| **MALAT1_43_176/177_3'3'_GC** | TAACATAGTTCAACCCACCATTTCTGGATTGTCCTTCCAGTTTCACCCTTCTCCTGGGTG |
| **MALAT1_44_176/177_3'3'_GC** | TTGCAGGCAAATTAATGGCCTTTCTGGATTGTCCTTCCAGTTTCACCCTTCTCCTGGGTG |
| **MALAT1_45_176/177_3'3'_GC** | TGTGGTTTTAATACCCTTCTTTTCTGGATTGTCCTTCCAGTTTCACCCTTCTCCTGGGTG |
| **MALAT1_46_176/177_3'3'_GC** | TTTCTCCACTTACTGGTTTATTTCTGGATTGTCCTTCCAGTTTCACCCTTCTCCTGGGTG |
| **MALAT1_47_176/177_3'3'_GC** | CTATCTTCTCCAGTCTACAATTTCTGGATTGTCCTTCCAGTTTCACCCTTCTCCTGGGTG |
| **MALAT1_48_176/177_3'3'_GC** | CAAGTCTGTTATGTTCACCTTTTCTGGATTGTCCTTCCAGTTTCACCCTTCTCCTGGGTG |

**Table S8.** MALAT 1 double label probe set.

| Probe | Sequence |
|---|---|
| **MALAT1_1_176/177_3'3'_GG** | GCCCTTCTATTGGTATTAATTTTCTGGATTGTCCTTCCAGTTTCACCCTTGTCCTGGGTG |
| **MALAT1_2_176/177_3'3'_GG** | CTCCCAATTAATCTTTCCATTTTCTGGATTGTCCTTCCAGTTTCACCCTTGTCCTGGGTG |
| **MALAT1_3_176/177_3'3'_GG** | TCTCCAAATTGTTTCATCCTTTTCTGGATTGTCCTTCCAGTTTCACCCTTGTCCTGGGTG |
| **MALAT1_4_176/177_3'3'_GG** | TTAGCTTTTTGTTTCCTAGCTTTCTGGATTGTCCTTCCAGTTTCACCCTTGTCCTGGGTG |
| **MALAT1_5_176/177_3'3'_GG** | ATCTTCTCAAGCTTTACCTTTTTCTGGATTGTCCTTCCAGTTTCACCCTTGTCCTGGGTG |
| **MALAT1_6_176/177_3'3'_GG** | ACTATATTTAAGGCCTTCCATTTCTGGATTGTCCTTCCAGTTTCACCCTTGTCCTGGGTG |
| **MALAT1_7_176/177_3'3'_GG** | TACTTCCGTTACGAAAGTCCTTTCTGGATTGTCCTTCCAGTTTCACCCTTGTCCTGGGTG |
| **MALAT1_8_176/177_3'3'_GG** | CTGGGTCAGCTGTCAATTAATTTCTGGATTGTCCTTCCAGTTTCACCCTTGTCCTGGGTG |
| **MALAT1_9_176/177_3'3'_GG** | TCAGTCCTAGCTTCATCAAATTTCTGGATTGTCCTTCCAGTTTCACCCTTGTCCTGGGTG |
| **MALAT1_10_176/177_3'3'_GG** | AACAACATATTGCCGACCTCTTTCTGGATTGTCCTTCCAGTTTCACCCTTGTCCTGGGTG |
| **MALAT1_11_176/177_3'3'_GG** | AGTCATTTGCCTTTAGGATTTTTCTGGATTGTCCTTCCAGTTTCACCCTTGTCCTGGGTG |
| **MALAT1_12_176/177_3'3'_GG** | AACTGTAAACCTGTGGTGGTTTTCTGGATTGTCCTTCCAGTTTCACCCTTGTCCTGGGTG |
| **MALAT1_13_176/177_3'3'_GG** | CTCAACGTGAGAACTGCTCTTTTCTGGATTGTCCTTCCAGTTTCACCCTTGTCCTGGGTG |
| **MALAT1_14_176/177_3'3'_GG** | CTCTAACCCAGTTTGTCAATTTTCTGGATTGTCCTTCCAGTTTCACCCTTGTCCTGGGTG |
| **MALAT1_15_176/177_3'3'_GG** | CCAAGGATAAAAGCAGCTCCTTTCTGGATTGTCCTTCCAGTTTCACCCTTGTCCTGGGTG |
| **MALAT1_16_176/177_3'3'_GG** | TGAACCAAAGCTGCACTGTGTTTCTGGATTGTCCTTCCAGTTTCACCCTTGTCCTGGGTG |
| **MALAT1_17_176/177_3'3'_GG** | ACTGCCAACTAATTGCCAATTTTCTGGATTGTCCTTCCAGTTTCACCCTTGTCCTGGGTG |
| **MALAT1_18_176/177_3'3'_GG** | CCAGTGGCTCATATTTAACTTTTCTGGATTGTCCTTCCAGTTTCACCCTTGTCCTGGGTG |
| **MALAT1_19_176/177_3'3'_GG** | ACTTTCCTTGCCCAAATTAATTTCTGGATTGTCCTTCCAGTTTCACCCTTGTCCTGGGTG |
| **MALAT1_20_176/177_3'3'_GG** | CCCAATGGAGGTATGACATATTTCTGGATTGTCCTTCCAGTTTCACCCTTGTCCTGGGTG |
| **MALAT1_21_176/177_3'3'_GG** | CATGCAATACTGCAGATGCATTTCTGGATTGTCCTTCCAGTTTCACCCTTGTCCTGGGTG |
| **MALAT1_22_176/177_3'3'_GG** | AGGAGAAAGTGCCATGGTTGTTTCTGGATTGTCCTTCCAGTTTCACCCTTGTCCTGGGTG |
| **MALAT1_23_176/177_3'3'_GG** | TTTCTCAATCCTGAAATCCCTTTCTGGATTGTCCTTCCAGTTTCACCCTTGTCCTGGGTG |
| **MALAT1_24_176/177_3'3'_GG** | TGAAGTGTACTATCCCATCATTTCTGGATTGTCCTTCCAGTTTCACCCTTGTCCTGGGTG |
| **MALAT1_25_176/177_3'3'_GG** | TCCTGATCTGGTCCATTAAATTTCTGGATTGTCCTTCCAGTTTCACCCTTGTCCTGGGTG |
| **MALAT1_26_176/177_3'3'_GG** | TCTTTCCTGCCTTAAAGTTATTTCTGGATTGTCCTTCCAGTTTCACCCTTGTCCTGGGTG |
| **MALAT1_27_176/177_3'3'_GG** | TGTCAATTTATAGACCCCTGTTTCTGGATTGTCCTTCCAGTTTCACCCTTGTCCTGGGTG |
| **MALAT1_28_176/177_3'3'_GG** | AAACATTGCCTACCACTCTATTTCTGGATTGTCCTTCCAGTTTCACCCTTGTCCTGGGTG |
| **MALAT1_29_176/177_3'3'_GG** | CCTGAATGGCTTCATGAAGGTTTCTGGATTGTCCTTCCAGTTTCACCCTTGTCCTGGGTG |
| **MALAT1_30_176/177_3'3'_GG** | TGCATTTACTTGCCAACAGATTTCTGGATTGTCCTTCCAGTTTCACCCTTGTCCTGGGTG |
| **MALAT1_31_176/177_3'3'_GG** | GTCGTTTCACAATGCATTCTTTTCTGGATTGTCCTTCCAGTTTCACCCTTGTCCTGGGTG |
| **MALAT1_32_176/177_3'3'_GG** | CAACACTCAGCCTTTATCACTTTCTGGATTGTCCTTCCAGTTTCACCCTTGTCCTGGGTG |
| **MALAT1_33_176/177_3'3'_GG** | TTTTTTCTTACTGGGTCTGGTTTCTGGATTGTCCTTCCAGTTTCACCCTTGTCCTGGGTG |
| **MALAT1_34_176/177_3'3'_GG** | ACTGGAAGCTCCTTCTATAGTTTCTGGATTGTCCTTCCAGTTTCACCCTTGTCCTGGGTG |
| **MALAT1_35_176/177_3'3'_GG** | TCATTTTGTCCACTGGTGAATTTCTGGATTGTCCTTCCAGTTTCACCCTTGTCCTGGGTG |
| **MALAT1_36_176/177_3'3'_GG** | TTGTCCCATAACTGATCTGATTTCTGGATTGTCCTTCCAGTTTCACCCTTGTCCTGGGTG |
| **MALAT1_37_176/177_3'3'_GG** | AACACAGTTTGCTCACATGCTTTCTGGATTGTCCTTCCAGTTTCACCCTTGTCCTGGGTG |
| **MALAT1_38_176/177_3'3'_GG** | TGACACTTCTCTTGACCTTATTTCTGGATTGTCCTTCCAGTTTCACCCTTGTCCTGGGTG |
| **MALAT1_39_176/177_3'3'_GG** | CACTCCAGAAAGAGGGAGTTTTTCTGGATTGTCCTTCCAGTTTCACCCTTGTCCTGGGTG |
| **MALAT1_40_176/177_3'3'_GG** | AGATCAAAAGGCACGGGGTGTTTCTGGATTGTCCTTCCAGTTTCACCCTTGTCCTGGGTG |
| **MALAT1_41_176/177_3'3'_GG** | ACAAGGATCCAAGCTACTGGTTTCTGGATTGTCCTTCCAGTTTCACCCTTGTCCTGGGTG |
| **MALAT1_42_176/177_3'3'_GG** | CATCGTTACCTTGAAACCGATTTCTGGATTGTCCTTCCAGTTTCACCCTTGTCCTGGGTG |
| **MALAT1_43_176/177_3'3'_GG** | TAACATAGTTCAACCCACCATTTCTGGATTGTCCTTCCAGTTTCACCCTTGTCCTGGGTG |
| **MALAT1_44_176/177_3'3'_GG** | TTGCAGGCAAATTAATGGCCTTTCTGGATTGTCCTTCCAGTTTCACCCTTGTCCTGGGTG |
| **MALAT1_45_176/177_3'3'_GG** | TGTGGTTTTAATACCCTTCTTTTCTGGATTGTCCTTCCAGTTTCACCCTTGTCCTGGGTG |
| **MALAT1_46_176/177_3'3'_GG** | TTTCTCCACTTACTGGTTTATTTCTGGATTGTCCTTCCAGTTTCACCCTTGTCCTGGGTG |
| **MALAT1_47_176/177_3'3'_GG** | CTATCTTCTCCAGTCTACAATTTCTGGATTGTCCTTCCAGTTTCACCCTTGTCCTGGGTG |
| **MALAT1_48_176/177_3'3'_GG** | CAAGTCTGTTATGTTCACCTTTTCTGGATTGTCCTTCCAGTTTCACCCTTGTCCTGGGTG |

### 11. **Supplementary Figures**

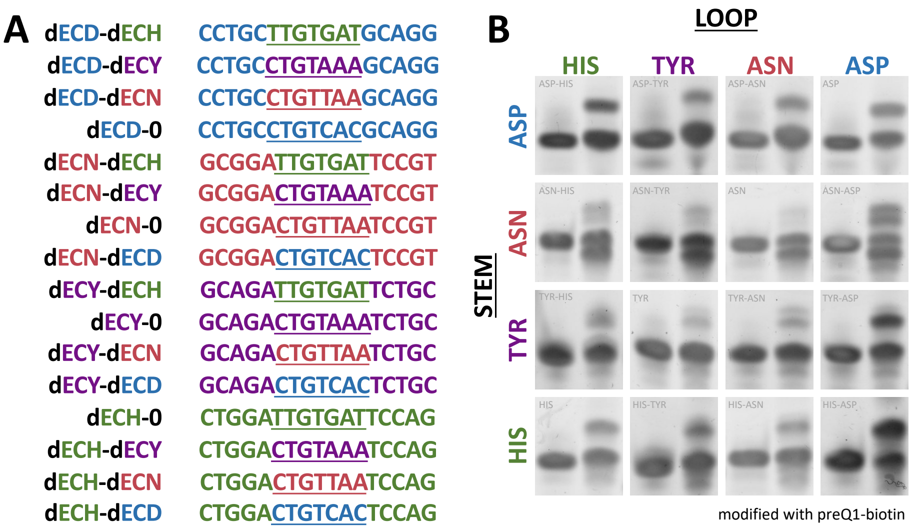


**Figure S1.** TGT labeling of DNA analog chimeras of queuine cognate tRNA anticodon loops. **(A)** Hairpin Sequences. **(B)** Urea PAGE gel shift analysis. For gel data, unmodified oligos are in the left lane and reaction products are in the right lane. Modification is determined by an upward gel shift of the oligo after insertion of preQ1-biotin.

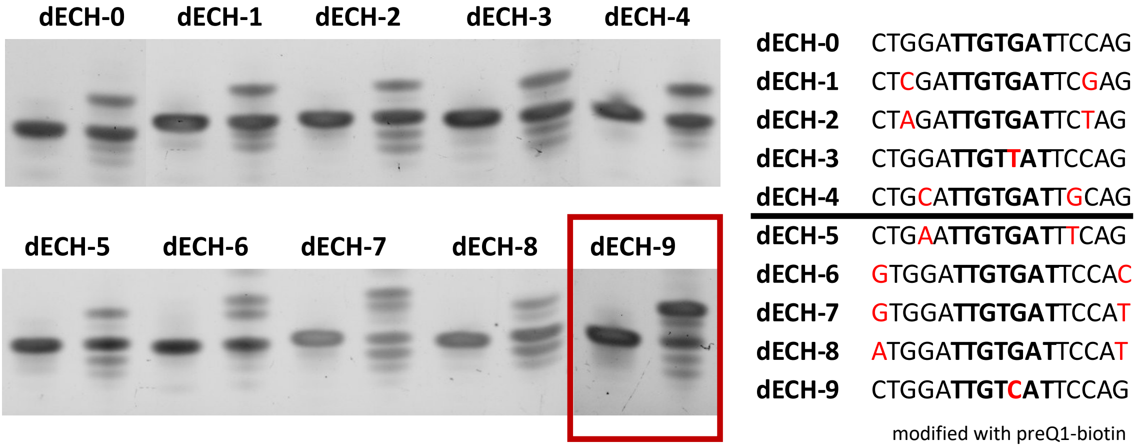


**Figure S2.** TGT modification of histidine hairpin mutants dECH1-9 with preQ1-biotin. For gel data, unmodified oligos are in the left lane and reaction products are in the right lane. Modification is determined by an upward gel shift of the oligo after insertion of preQ1-biotin.

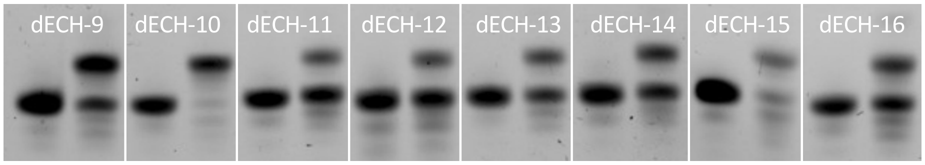


| | | | |
|---|---|---|---|
| dECH-9 | CTGGATTGTCATTCCAG | dECH-13 | CTGGATTGTCTTTCCAG |
| dECH-10 | CTGGATTGTCCTTCCAG | dECH-14 | CTGGATTGTCGTTCCAG |
| dECH-11 | CTGGATTGTTTTTCCAG | dECH-15 | CTGGATTGTTGTTCCAG |
| dECH-12 | CTGGATTGTTATTCCAG | dECH-16 | CTGGATTGTTCTTCCAG |

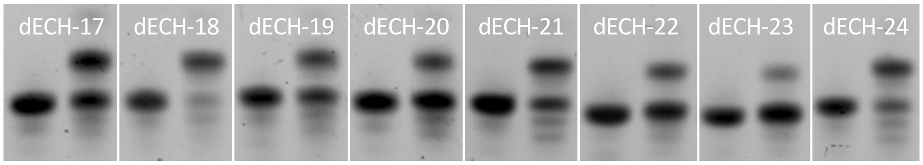


| | | | |
|---|---|---|---|
| dECH-17 | CTGGACTGTCACTCCAG | dECH-21 | CTGGACTGTCTCTCCAG |
| dECH-18 | CTGGACTGTCCCTCCAG | dECH-22 | CTGGACTGTCGCTCCAG |
| dECH-19 | CTGGACTGTTTCTCCAG | dECH-23 | CTGGACTGTTGCTCCAG |
| dECH-20 | CTGGACTGTTACTCCAG | dECH-24 | CTGGACTGTTCCTCCAG |

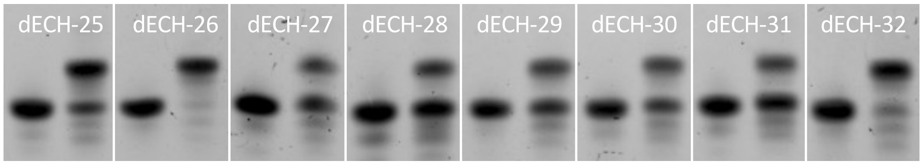


| | | | |
|---|---|---|---|
| dECH-25 | CTGGATTGTCACTCCAG | dECH-29 | CTGGATTGTCTCTCCAG |
| dECH-26 | CTGGATTGTCCCTCCAG | dECH-30 | CTGGATTGTCGCTCCAG |
| dECH-27 | CTGGATTGTTTCTCCAG | dECH-31 | CTGGATTGTTGCTCCAG |
| dECH-28 | CTGGATTGTTACTCCAG | dECH-32 | CTGGATTGTTCCTCCAG |

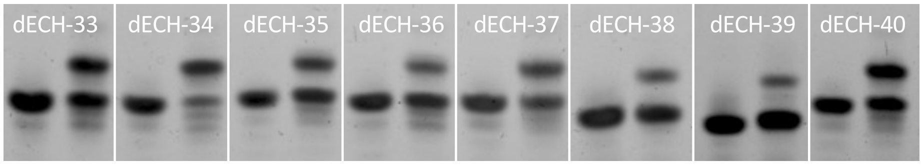


| | | | |
|---|---|---|---|
| dECH-33 | CTGGACTGTCATTCCAG | dECH-37 | CTGGACTGTCTTTCCAG |
| dECH-34 | CTGGACTGTCCTTCCAG | dECH-38 | CTGGACTGTCGTTCCAG |
| dECH-35 | CTGGACTGTTTTTCCAG | dECH-39 | CTGGACTGTTGTTCCAG |
| dECH-36 | CTGGACTGTTATTCCAG | dECH-40 | CTGGACTGTTCTTCCAG |

**Figure S3.** TGT modification of histidine hairpin loop mutants dECH9-40. Mutations are indicated in red. For gel data, unmodified oligos are in the left lane and reaction products are in the right lane. Modification is determined by an upward gel shift of the oligo after insertion of preQ1-biotin. Highlighted sequences are labeled with greater than 85% efficiency as determined by densitometry. The minimal loop sequence requirement is YTGTCCY or YTGTYCC where Y is a smaller pyrimidine nucleobase, C or T.

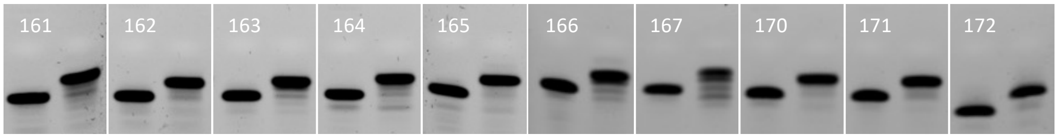


| | | | |
|---|---|---|---|
| 161_ala1_dECH-10 | cttgcTTGTCCTgcaag | 166_arg4_dECH-10 | acgacTTGTCCTgtcgt |
| 162_ala2_dECH-10 | cctgcTTGTCCTgcagg | 167_asn1_dECH-10 | gcggaTTGTCCTtccgt |
| 163_arg1_dECH-10 | agcccTTGTCCTgggct | 170_gln1_dECH-10 | ccggaTTGTCCTtccgg |
| 164_arg2_dECH-10 | ctcggTTGTCCTccgag | 171_gln2_dECH-10 | ccggtTTGTCCTaccgg |
| 165_arg3_dECH-10 | ctgccTTGTCCTggcag | 172_glu1_dECH-10 | ccgccTTGTCCTggcgg |

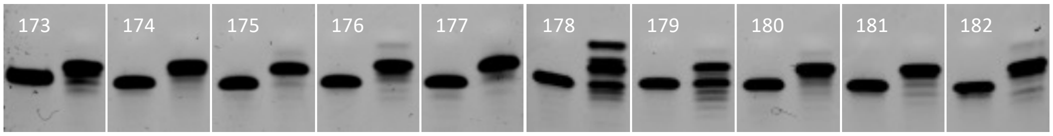


| | | | |
|---|---|---|---|
| 173_gly1_dECH-10 | agagcTTGTCCTgctct | 178_leu1_dECH-10 | gttgaTTGTCCTtcaac |
| 174_gly2_dECH-10 | tcagcTTGTCCTgctga | 179_leu2_dECH-10 | ctagcTTGTCCTgttag |
| 175_gly3_dECH-10 | cgaccTTGTCCTggtcg | 180_leu3_dECH-10 | ctaccTTGTCCTggtag |
| 176_his1_dECH-10 | ctggaTTGTCCTtccag | 181_leu4_dECH-10 | ccagaTTGTCCTtctgg |
| 177_ile1_dECH-10 | cacccTTGTCCTgggtg | 182_leu5_dECH-10 | agggaTTGTCCTtccct |

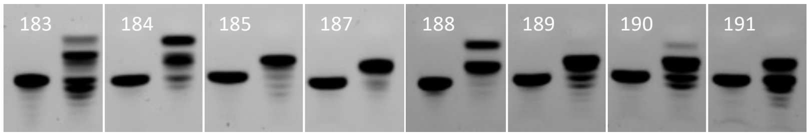


| | | | |
|---|---|---|---|
| 183_lys1_dECH-10 | gttgaTTGTCCTtcaat | 188_phe1_dECH-10 | ggggaTTGTCCTtcccc |
| 184_met1_dECH-10 | ggcgaTTGTCCTtcgct | 189_pro1_dECH-10 | actggTTGTCCTccagt |
| 185_met2_dECH-10 | catcaTTGTCCTtgatg | 190_pro2_dECH-10 | ccgtcTTGTCCTgtcgg |
| 187_met4_dECH-10 | tcgggTTGTCCTcccga | 191_pro3_dECH-10 | cttcgTTGTCCTcgagg |

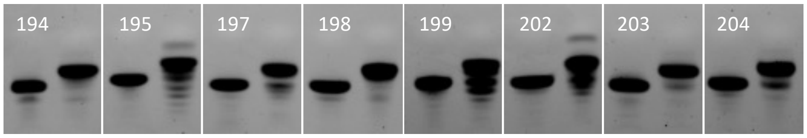


| | | | |
|---|---|---|---|
| 194_ser3_dECH-10 | ctcccTTGTCCTgggag | 199_thr4_dECH-10 | actgaTTGTCCTtcagt |
| 195_ser4_dECH-10 | cacgcTTGTCCTgtgtg | 202_tyr1_dECH-10 | gcagaTTGTCCTtctgc |
| 197_thr2_dECH-10 | tcgcaTTGTCCTtgcga | 203_val1_dECH-10 | ccaccTTGTCCTggtgg |
| 198_thr3_dECH-10 | cgcatTTGTCCTatgcg | 204_val2_dECH-10 | cctccTTGTCCTggagg |

**Figure S4.** TGT modification of anticodon stem/dECH-10 loop chimeras. Anticodon stems of all tRNAs were appended with the dECH-10 loop to determine the importance of stem vs loop sequence for TGT labeling. For gel data, unmodified oligos are in the left lane and reaction products are in the right lane. Modification is determined by an upward gel shift of the oligo after insertion of preQ1-biotin. Redundant hairpins where two or more tRNA anticodon arms had identical stem sequences were omitted from this figure. See Table S3 for a complete list of anticodon stem/dECH-10 loop chimeras including redundant hairpins. Highlighted sequences are labeled with greater than 85% efficiency as determined by densitometry.

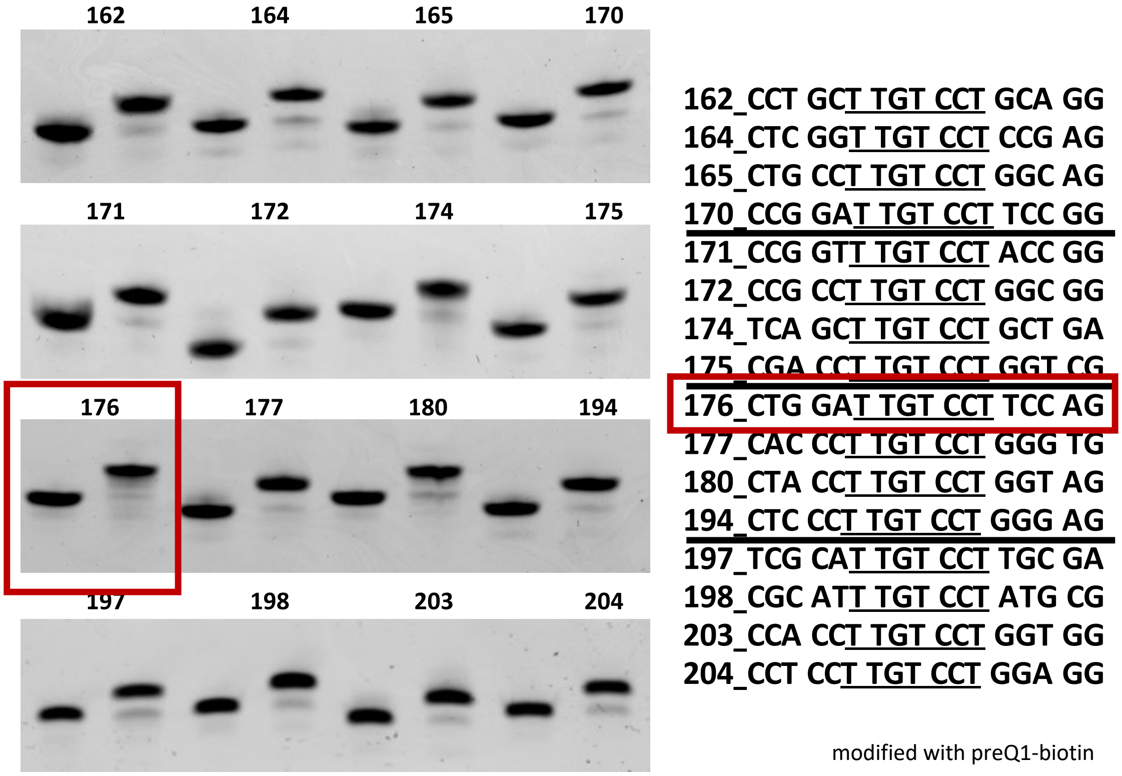


**Figure S5.** TGT modification of 16 distinct hairpin sequences. For gel data, unmodified oligos are in the left lane and reaction products are in the right lane. Modification is determined by an upward gel shift of the oligo after insertion of preQ1-biotin.

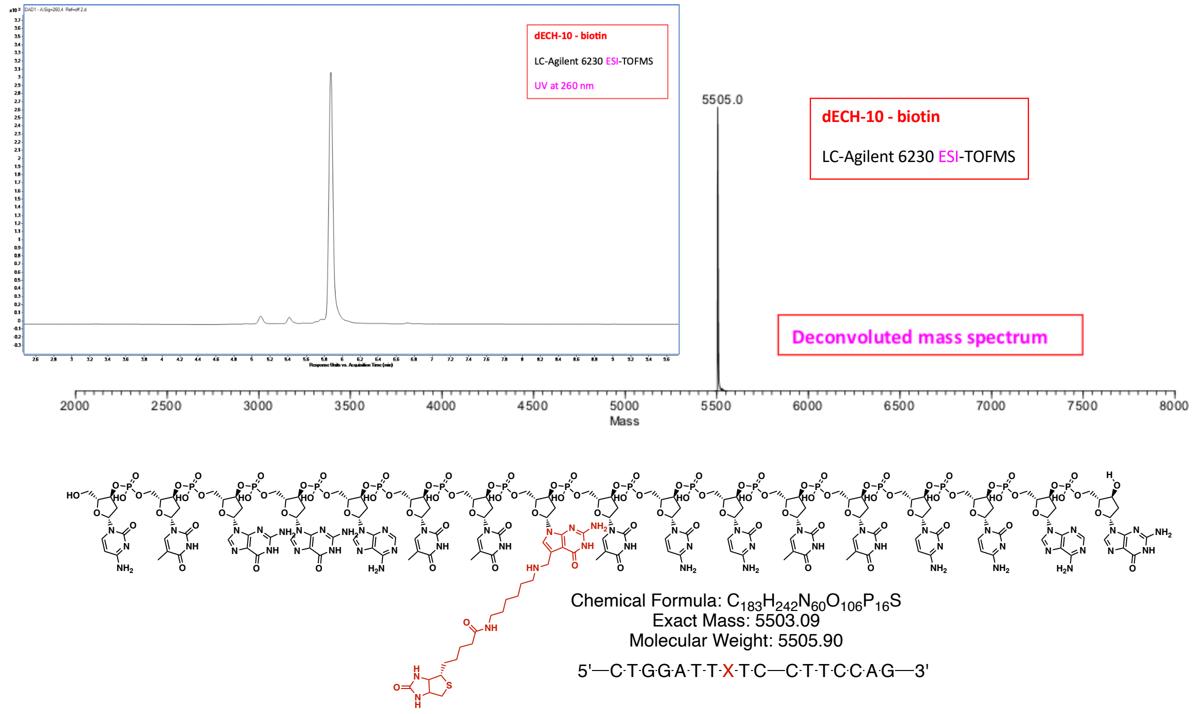


**Figure S6.** LCMS verification of dECH-10 modification with preQ1-biotin by *E. coli* TGT.

dECH-10 dECH-10 ΔC

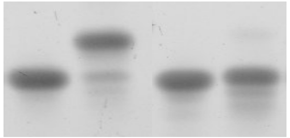

modified with PreQ1-biotin

dECH-10 CTGGA**TTGTCCT**TCCAG
dECH-10 ΔC CTGGA**TTCTCCT**TCCAG

**Figure S7.** Confirmation of specific guanine exchange. Mutation of the key G nucleobase to C results in no observable product. For gel data, unmodified oligos are in the left lane and reaction products are in the right lane. Modification is determined by an upward gel shift of the oligo after insertion of preQ1-biotin.

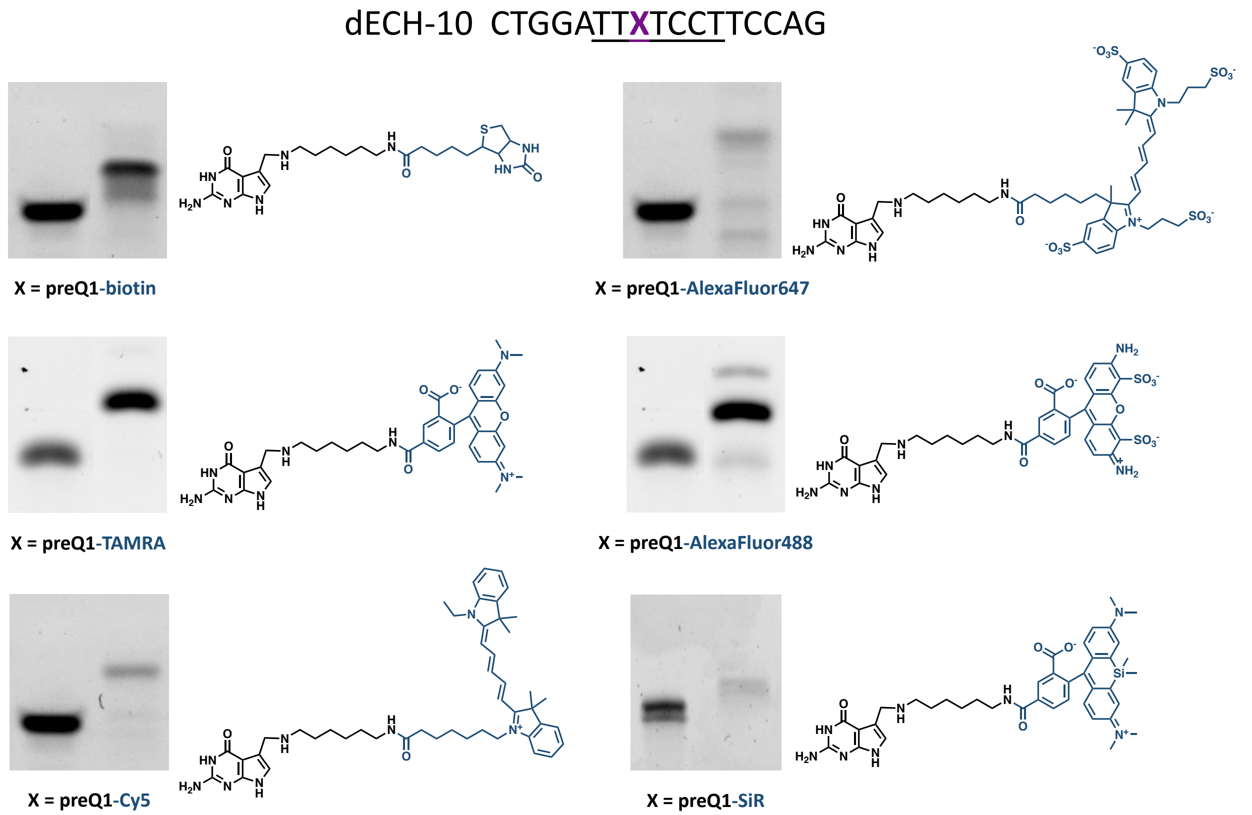


**Figure S8.** Substrate scope of DNA-TAG. For gel data, unmodified oligos are in the left lane and reaction products are in the right lane. Modification is determined by an upward gel shift of the oligo after insertion of of preQ1 probes. Cy5, AlexaFluor647, and silicon rhodamine (SiR) dyes are quenched by gel red making the product band faint. See section 12 for confirmation of labeling by MS.

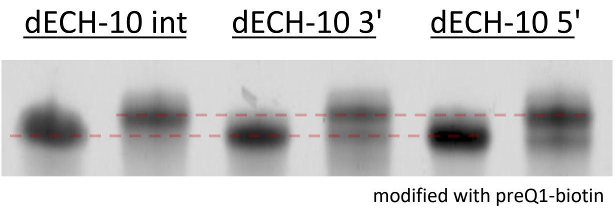


dECH-10 int ACCATCATTTTCATATCCTCCACTGGA**TTGTCCT**TCCAGACCACCATCATTTGCAATGA

dECH-10 3' ACCATCATTTTCATATCCTCCAACCACCATCATTTGCAATGACTGGA**TTGTCCT**TCCAG

dECH-10 5' CTGGA**TTGTCCT**TCCAGACCATCATTTTCATATCCTCCAACCACCATCATTTGCAATGA

**Figure S9.** The dECH-10 hairpin is labeled when inserted into the center, or appended to either end, of a DNA substrate. For gel data, unmodified oligos are in the left lane and reaction products are in the right lane. Modification is determined by an upward gel shift of the oligo after insertion of preQ1-biotin.

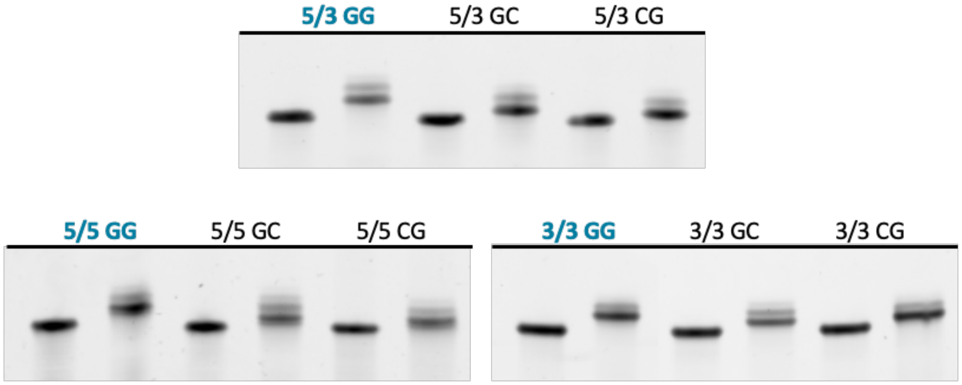


| oligonucleotide | sequence | expected m/z | deconvoluted m/z |
|---|---|---|---|
| **5/3_176/177_GG** | **ctggaTTgTCCTtccagCATCATTTTCATATCCTCCAcacccTTgTCCTgggtg** | **17098.02** | **17099** |
| 5/3_176/177_GC | ctggaTTgTCCTtccagCATCATTTTCATATCCTCCAcacccTTcTCCTgggtg | 16704.81 | 16705 |
| 5/3_176/177_CG | ctggaTTcTCCTtccagCATCATTTTCATATCCTCCAcacccTTgTCCTgggtg | 16704.81 | 16705 |
| **5/5_176/177_GG** | **ctggaTTgTCCTtccagTTTcacccTTgTCCTgggtgCATCATTTTCATATCCTCCA** | **18010.62** | **18012** |
| 5/5_176/177_GC | ctggaTTgTCCTtccagTTTcacccTTcTCCTgggtgCATCATTTTCATATCCTCCA | 17617.41 | 17617 |
| 5/5_176/177_CG | ctggaTTcTCCTtccagTTTcacccTTgTCCTgggtgCATCATTTTCATATCCTCCA | 17617.41 | 17617 |
| **3/3_176/177_GG** | **CATCATTTTCATATCCTCCActggaTTgTCCTtccagTTTcacccTTgTCCTgggtg** | **18010.62** | **18011** |
| 3/3_176/177_GC | CATCATTTTCATATCCTCCActggaTTgTCCTtccagTTTcacccTTcTCCTgggtg | 17617.41 | 17617 |
| 3/3_176/177_CG | CATCATTTTCATATCCTCCActggaTTcTCCTtccagTTTcacccTTgTCCTgggtg | 17617.41 | 17617 |

**Figure S10.** Single or double labeling of DNA substrates bearing orthogonal hairpins with preQ1-biotin. Single label constructs have a G→C mutation in one of the two recognition element hairpins. Modification is determined by an upward gel shift of the oligo after insertion of preQ1-biotin. For gel data, unmodified oligos are in the left lane and reaction products are in the right lane. Dual labeling was successful in tandem at either end, and when separated at both ends. Highlighted oligos indicate GG substrates.

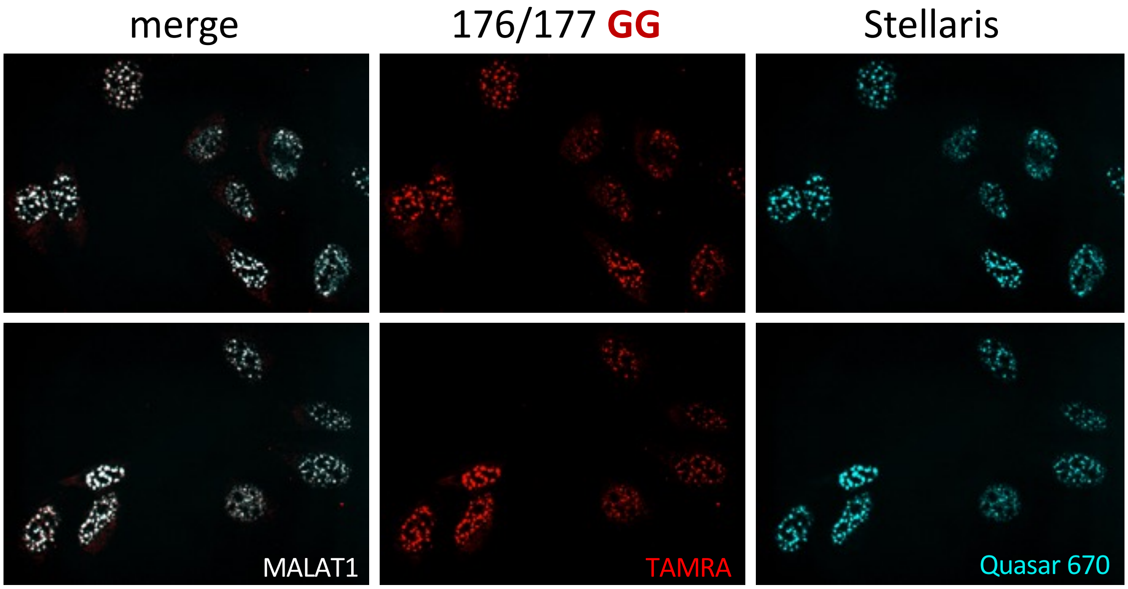


**Figure S11.** RNA FISH detection of MALAT1 with doubly labeled DNA-TAG generated probe set and Stellaris probe set. The DNA-TAG probe set reliably reports on MALAT1 as seen by colocalization with the commercially available probe set from Stellaris.

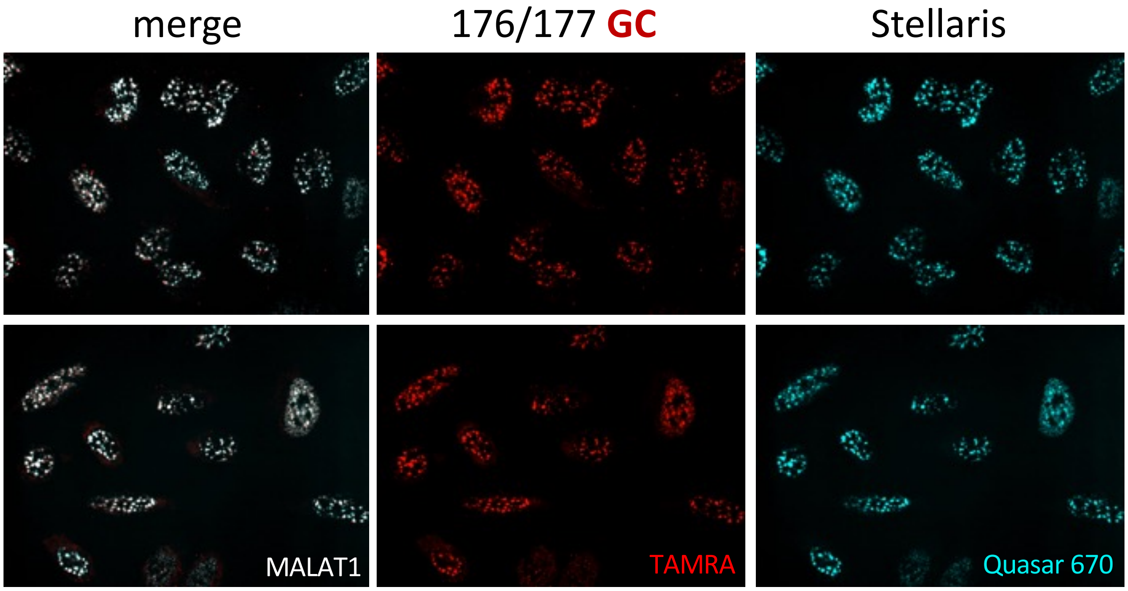


**Figure S12.** RNA FISH detection of MALAT1 with singly labeled DNA-TAG generated probe set and Stellaris probe set. The DNA-TAG probe set reliably reports on MALAT1 as seen by colocalization with the commercially available probe set from Stellaris.

### 12. MS of PreQ1 Probes

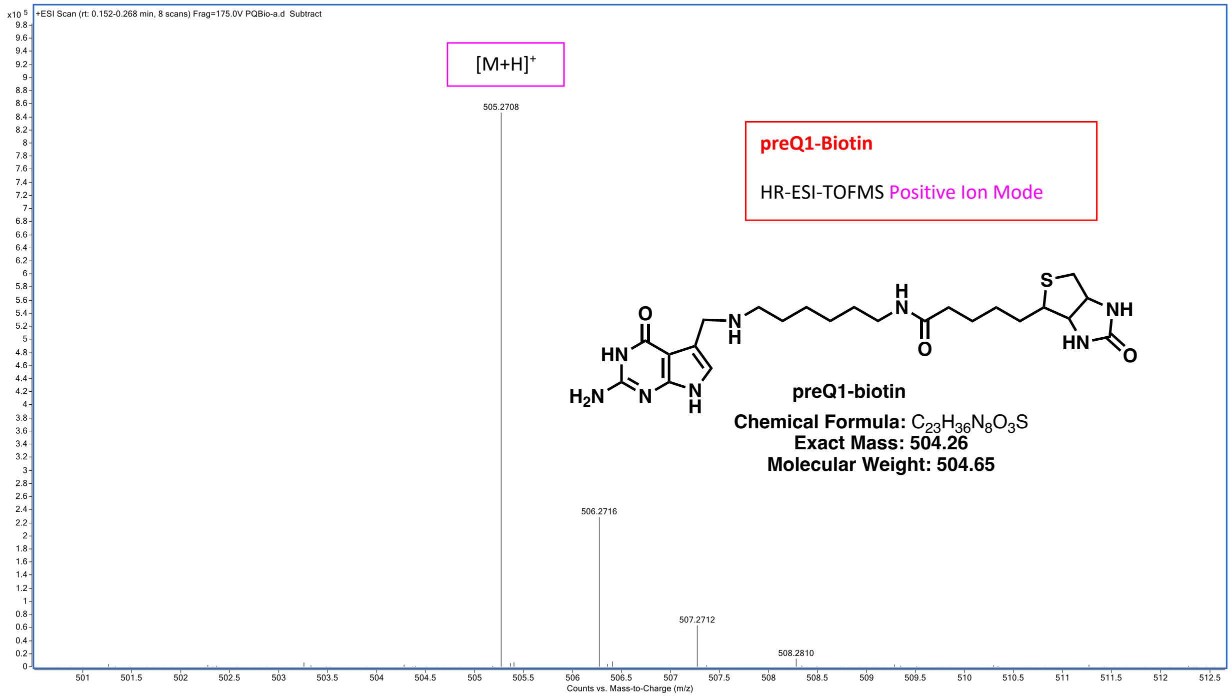


| Mass Measured | Theo. Mass | Delta (ppm) | Composition |
|---|---|---|---|
| 505.2708 | 505.2704 | 0.8 | $[C_{23}H_{37}N_8O_3S]^+$ |

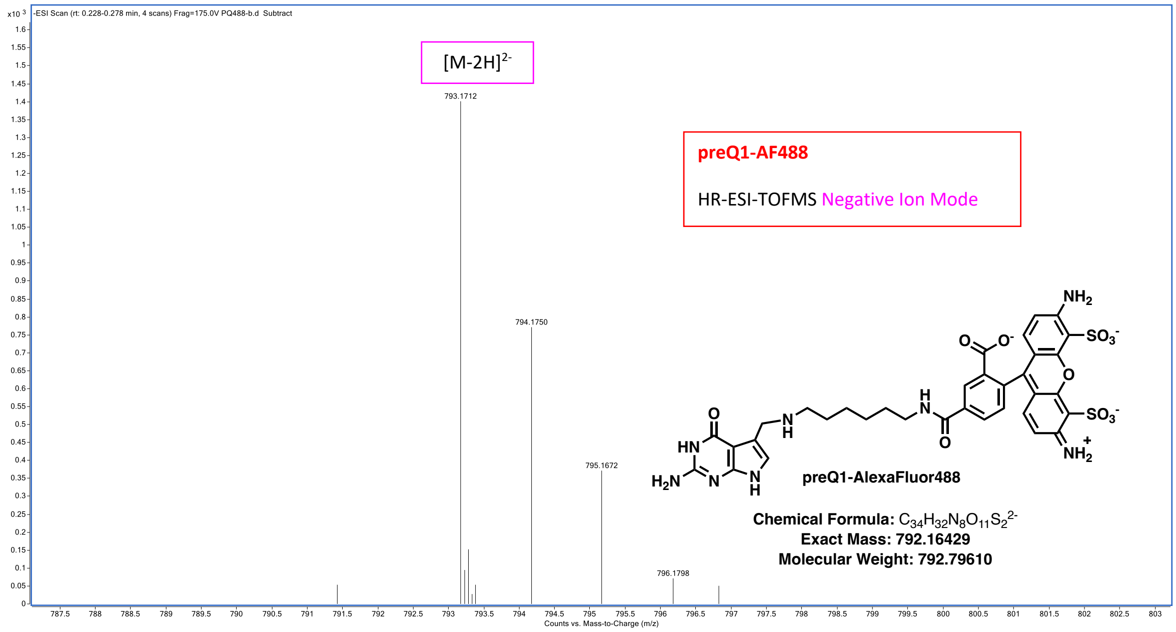


| Mass Measured | Theo. Mass | Delta (ppm) | Composition |
|---|---|---|---|
| 793.1712 | 793.1716 | -0.5 | $[C_{34}H_{32}N_8O_{11}S_2]^{2-}$ |

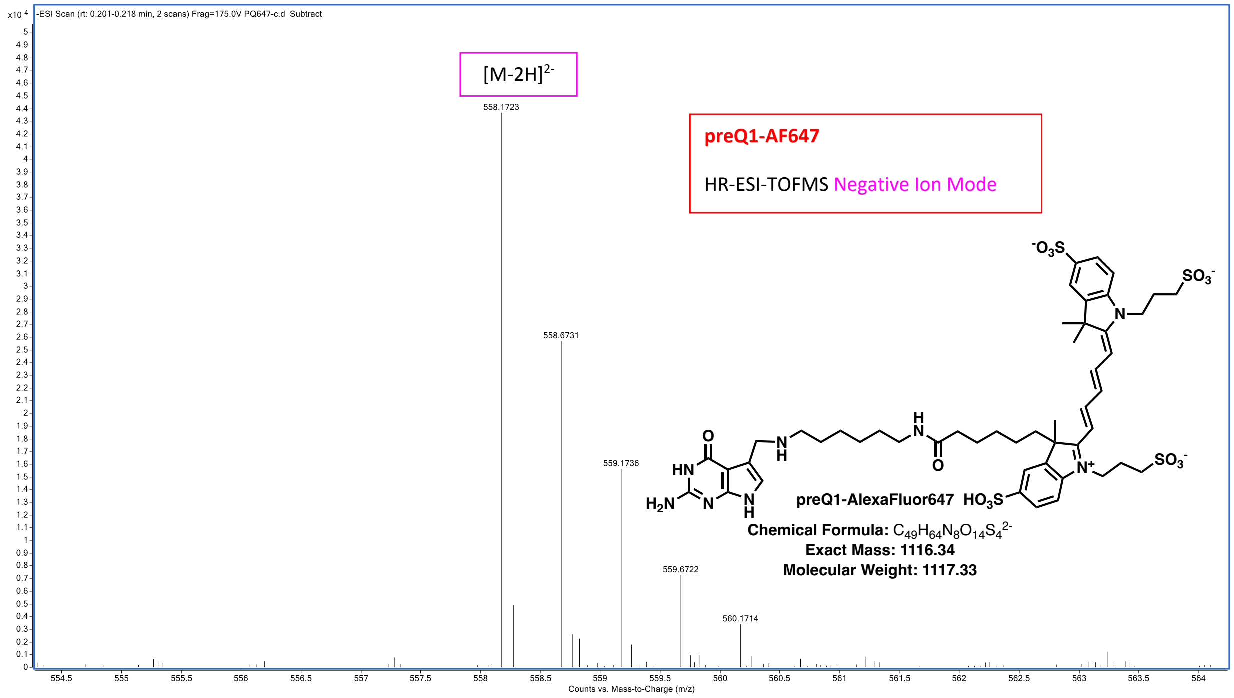


| Mass Measured | Theo. Mass | Delta (ppm) | Composition |
|---|---|---|---|
| 558.1723 | 558.1718 | -0.3 | $[C_{49}H_{64}N_8O_{14}S_4]^{2-}$ |

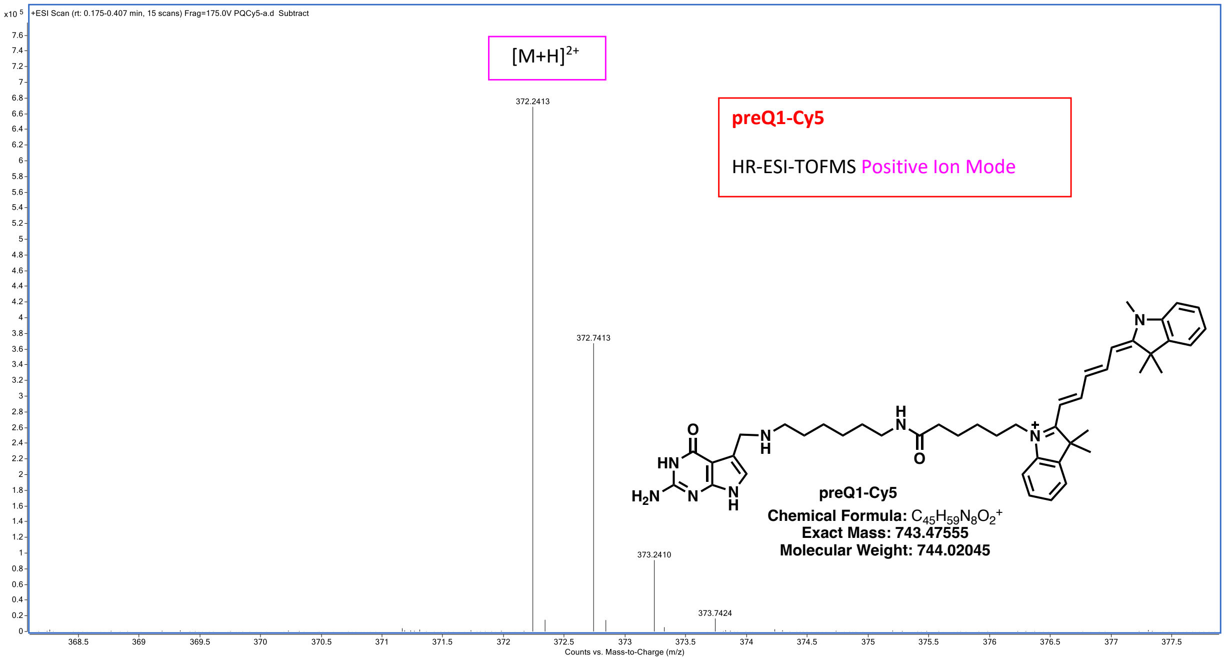


| Mass Measured | Theo. Mass | Delta (ppm) | Composition |
|---|---|---|---|
| 372.2413 | 372.2414 | -0.3 | $[C_{45}H_{60}N_8O_2]^{2+}$ |

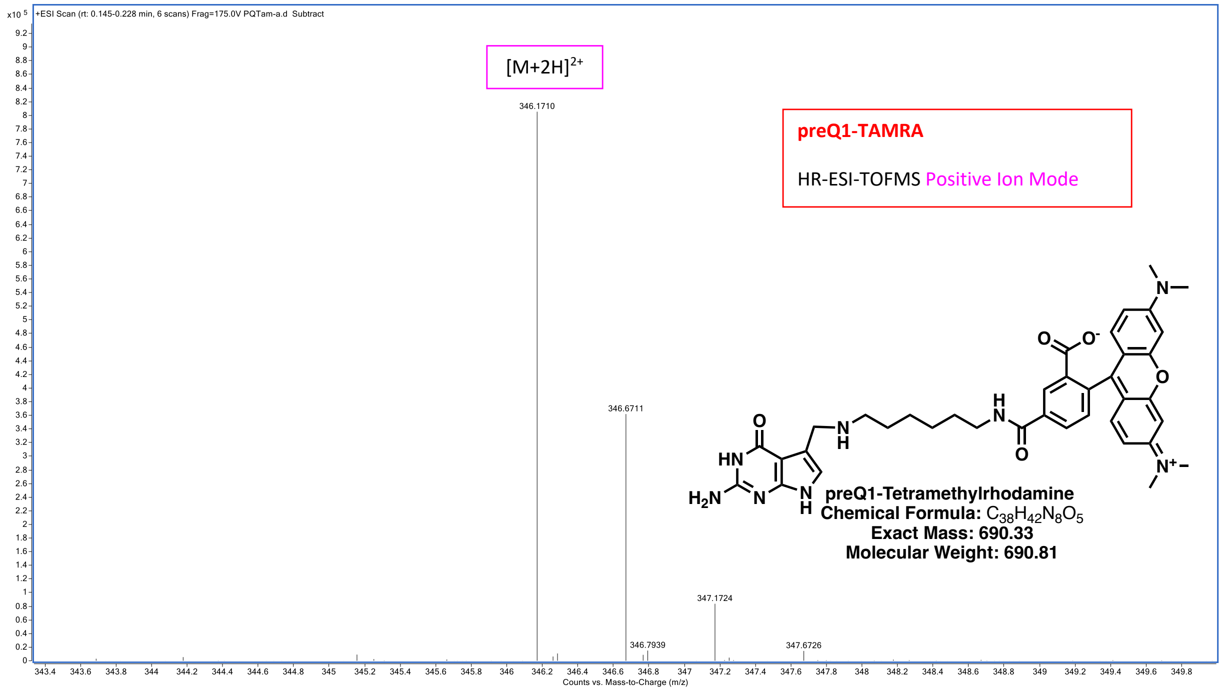


| Mass Measured | Theo. Mass | Delta (ppm) | Composition |
|---|---|---|---|
| 346.1710 | 346.1712 | -0.6 | $[C_{38}H_{44}N_8O_5]^{2+}$ |

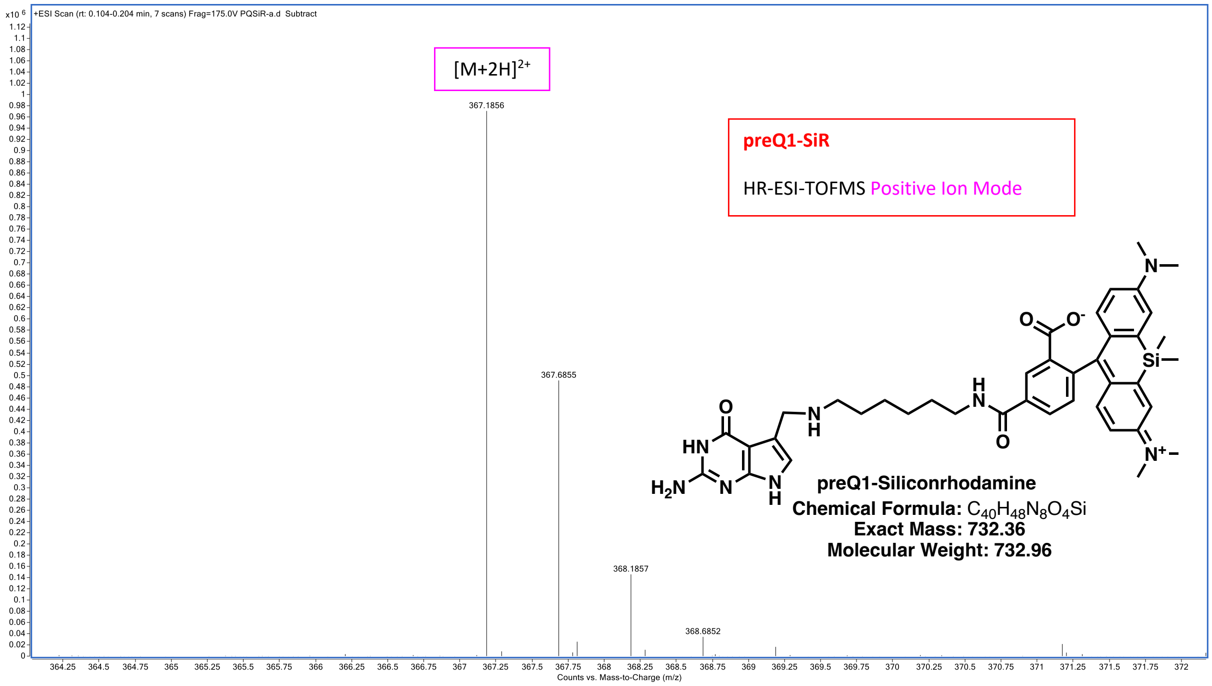


| Mass Measured | Theo. Mass | Delta (ppm) | Composition |
|---|---|---|---|
| 367.1856 | 367.1857 | -0.3 | $[C_{40}H_{50}N_8O_4Si]^{2+}$ |

### 13. MS of dECH-10 and modified oligos

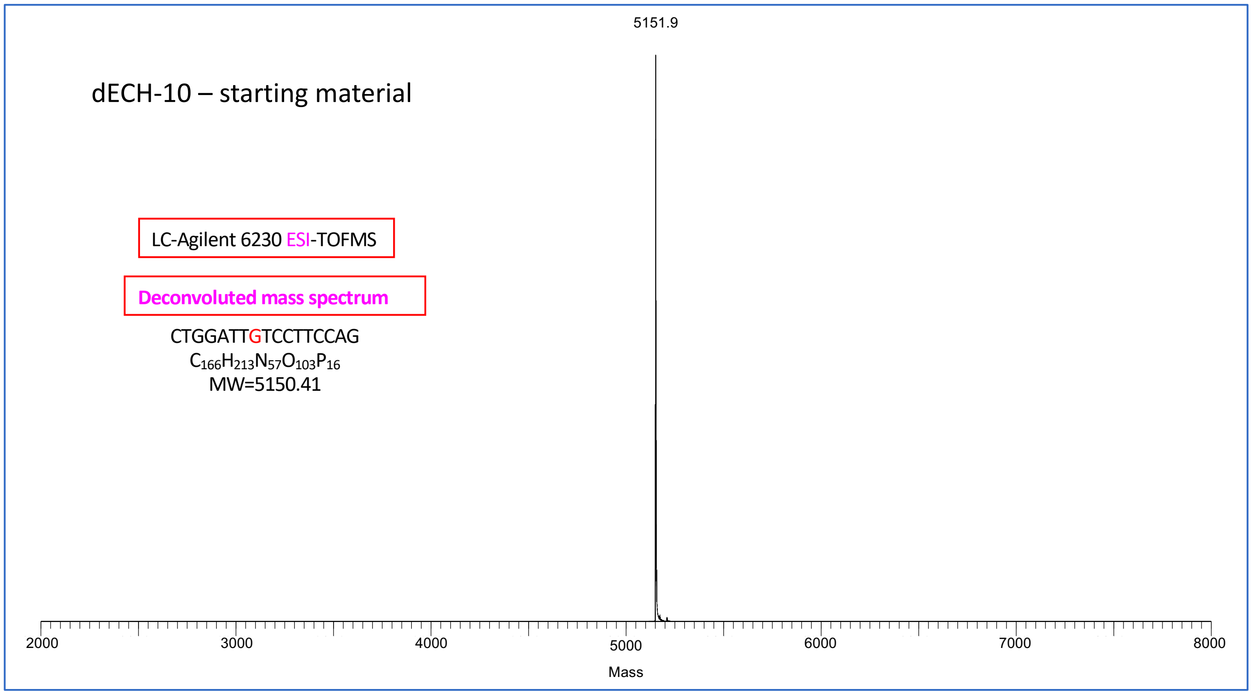


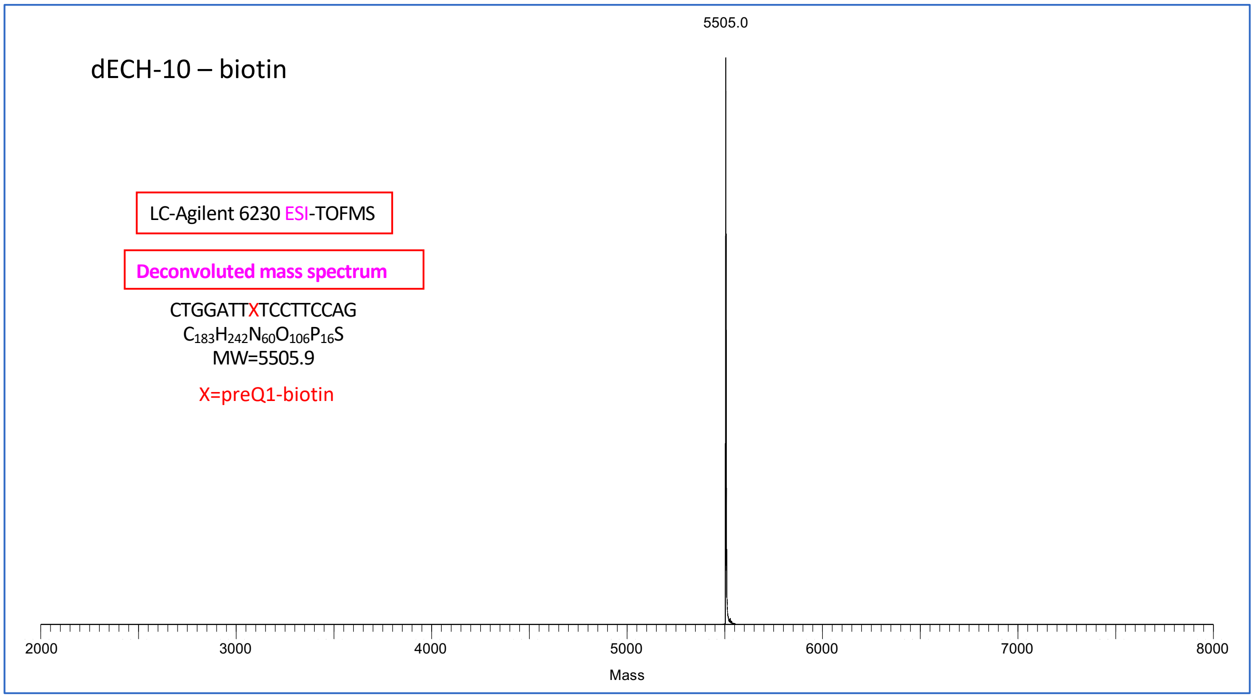

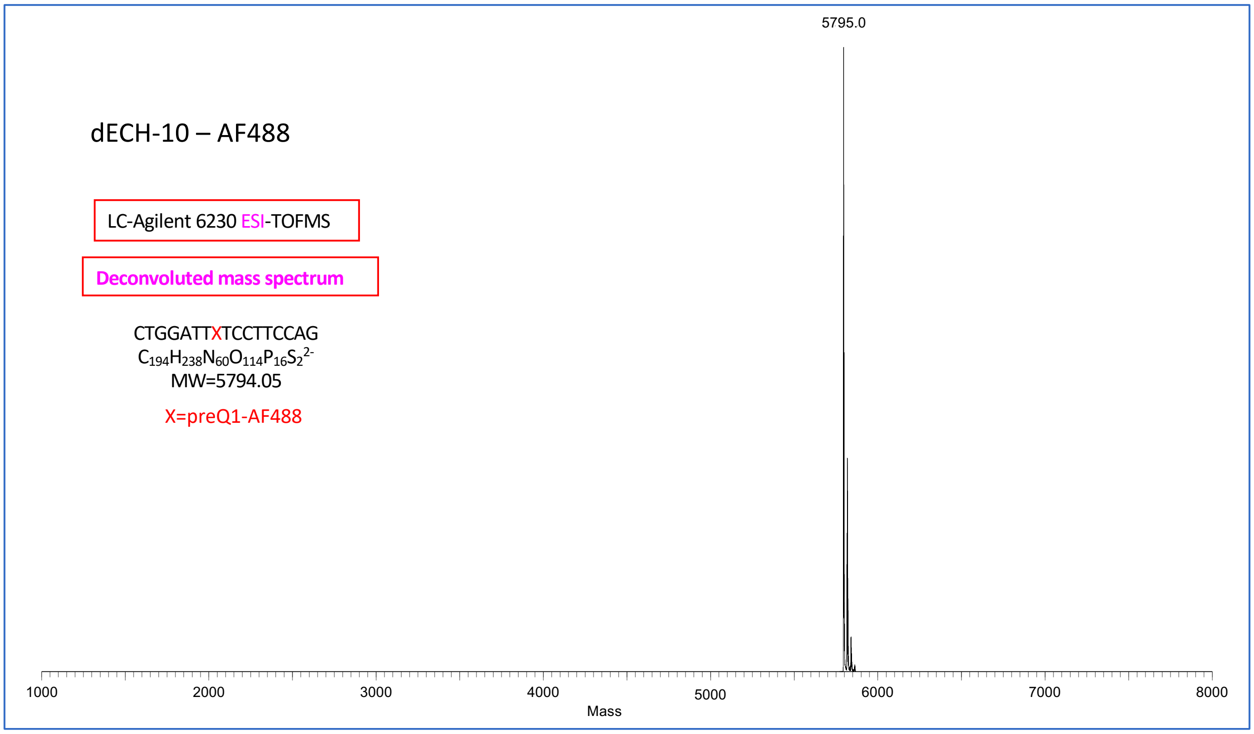
dECH-10 – AF488
LC-Agilent 6230 ESI-TOFMS
Deconvoluted mass spectrum
CTGGATTXTCCTTCCAG
$C_{194}H_{238}N_{60}O_{114}P_{16}S_2^{2-}$
MW=5794.05
X=preQ1-AF488
5795.0
1000
2000
3000
4000
5000
6000
7000
8000
Mass

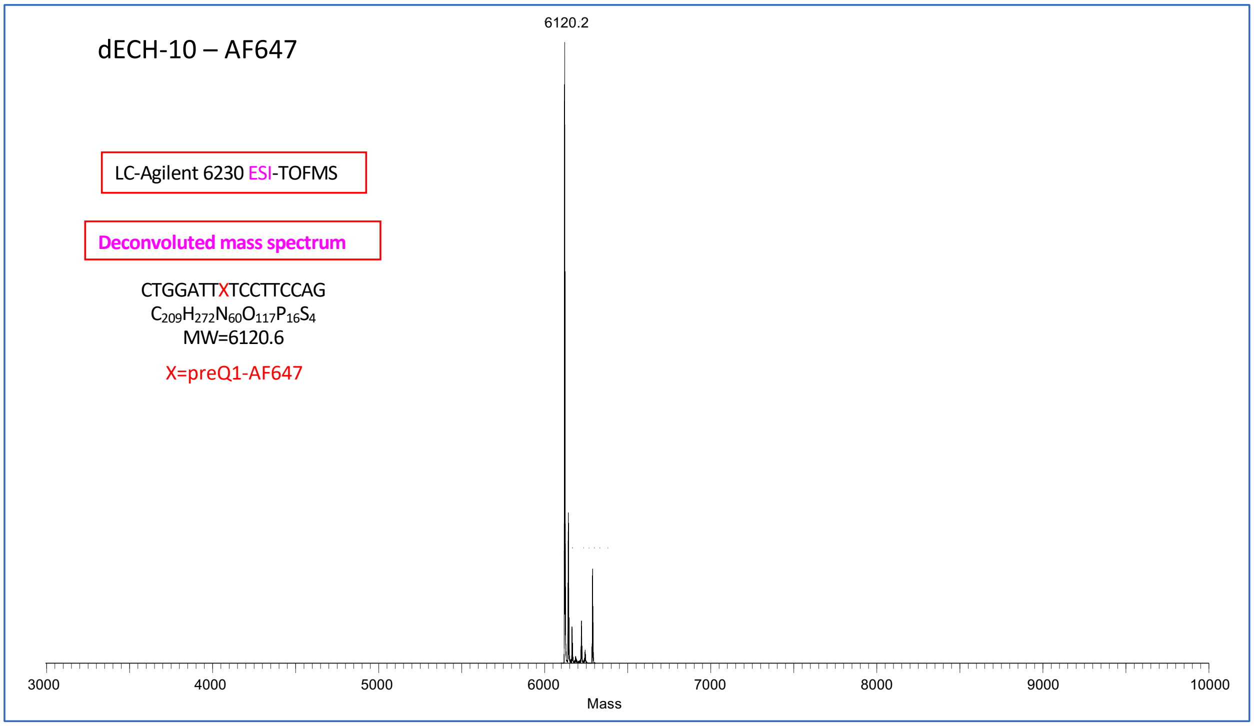
dECH-10 – AF647
LC-Agilent 6230 ESI-TOFMS
Deconvoluted mass spectrum
CTGGATTXTCCTTCCAG
$C_{209}H_{272}N_{60}O_{117}P_{16}S_4$
MW=6120.6
X=preQ1-AF647
6120.2
3000
4000
5000
6000
7000
8000
9000
10000
Mass

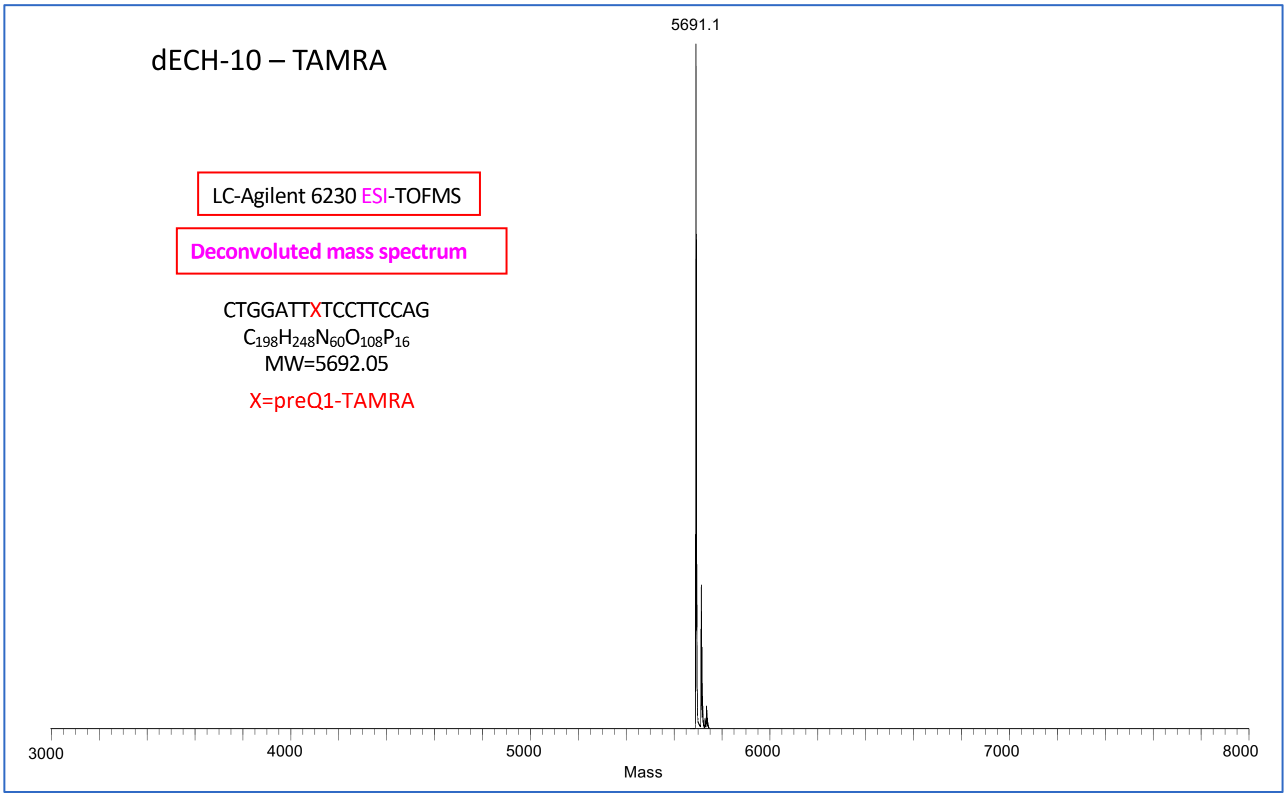
dECH-10 – TAMRA
5691.1
LC-Agilent 6230 ESI-TOFMS
Deconvoluted mass spectrum
CTGGATTXTCCTTCCAG
$C_{198}H_{248}N_{60}O_{108}P_{16}$
MW=5692.05
X=preQ1-TAMRA
3000
4000
5000
6000
7000
8000
Mass

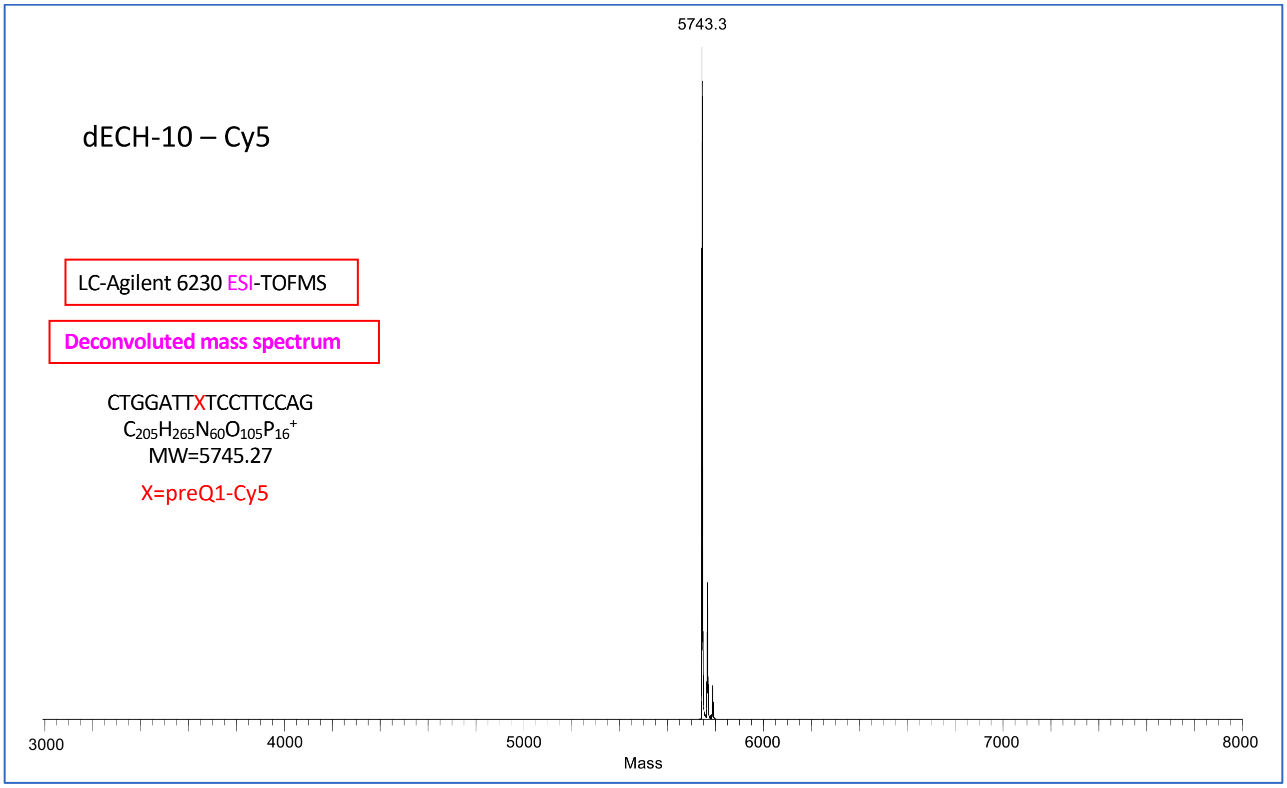
dECH-10 – Cy5
5743.3
LC-Agilent 6230 ESI-TOFMS
Deconvoluted mass spectrum
CTGGATTXTCCTTCCAG
$C_{205}H_{265}N_{60}O_{105}P_{16}^{+}$
MW=5745.27
X=preQ1-Cy5
3000
4000
5000
6000
7000
8000
Mass

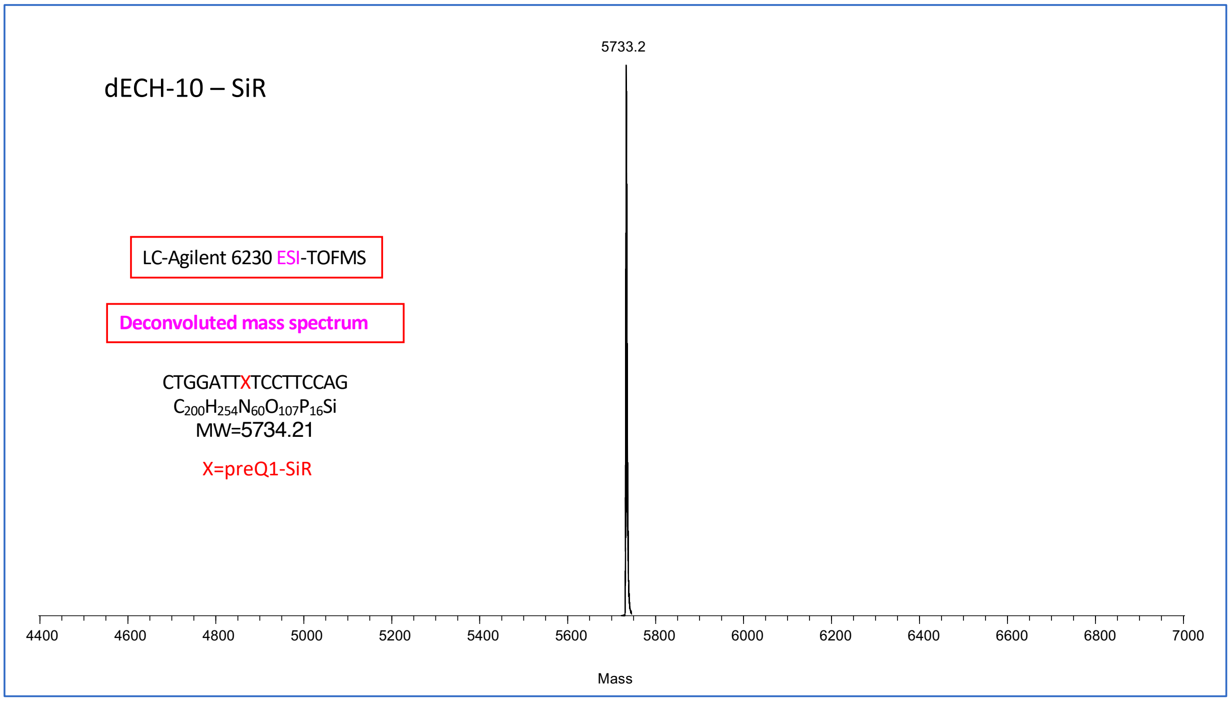
5733.2
dECH-10 – SiR
LC-Agilent 6230 ESI-TOFMS
Deconvoluted mass spectrum
CTGGATTXTCCTTCCAG
$C_{200}H_{254}N_{60}O_{107}P_{16}Si$
MW=5734.21
X=preQ1-SiR
4400
4600
4800
5000
5200
5400
5600
5800
6000
6200
6400
6600
6800
7000
Mass
